# Phage anti-CBASS protein simultaneously sequesters cyclic trinucleotides and dinucleotides

**DOI:** 10.1101/2023.06.01.543220

**Authors:** Xueli Cao, Yu Xiao, Erin Huiting, Xujun Cao, Dong Li, Jie Ren, Linlin Guan, Yu Wang, Lingyin Li, Joseph Bondy-Denomy, Yue Feng

**Author notes:** Authors contributed equally (Xueli Cao, Yu Xiao, Erin Huiting).

## Abstract

CBASS is a common anti-phage immune system that uses cyclic oligonucleotide signals to activate effectors and limit phage replication. In turn, phages encode anti-CBASS (Acb) proteins. We recently uncovered a widespread phage anti-CBASS protein Acb2 that acts as a “sponge” by forming a hexamer complex with three cGAMP molecules. Here, we identified that Acb2 binds and sequesters many CBASS and cGAS-produced cyclic dinucleotides *in vitro* and inhibits cGAMP-mediated STING activity in human cells. Surprisingly, Acb2 also binds CBASS cyclic trinucleotides 3’3’3’-cyclic AMP-AMP-AMP (cA3) and 3’3’3’-cAAG with high affinity. Structural characterization identified a distinct binding pocket within the Acb2 hexamer that binds two cyclic trinucleotide molecules and another binding pocket that binds to cyclic dinucleotides. Binding in one pocket does not allosterically alter the other, such that one Acb2 hexamer can simultaneously bind two cyclic trinucleotides and three cyclic dinucleotides. Phage-encoded Acb2 provides protection from Type III-C CBASS that uses cA3 signaling molecules *in vivo* and blocks cA3-mediated activation of the endonuclease effector *in vitro*. Altogether, Acb2 sequesters nearly all known CBASS signaling molecules through two distinct binding pockets and therefore serves as a broad-spectrum inhibitor of cGAS-based immunity.

## Introduction

Anti-viral immune pathways across all kingdoms of life sense and respond to viral infection. Cyclic GMP-AMP synthase (cGAS) is an evolutionarily conserved enzyme that performs a pivotal role in innate immunity against viruses (Li et al., 2013). In mammalian cells, cGAS binds viral DNA and is activated to produce 2’,3’-cyclic GMP-AMP (2’,3’-cGAMP) dinucleotides, which binds to and activates the STING (stimulator of interferon genes) effector protein to initiate a potent interferon response (Burdette et al., 2011; Ishikawa and Barber, 2008). In bacteria, cGAS-like enzymes named cGAS/DncV-like nucleotidyltransferases (CD-NTases) have been identified and enzymatically characterized (Burroughs et al., 2015; Davies et al., 2012; Whiteley et al., 2019). CD-NTases have been classified into 8 enzymatic clades and at least 12 cyclic di- and tri-nucleotide products have been identified (Burroughs et al., 2015; Davies et al., 2012; Duncan-Lowey et al., 2021; Fatma et al., 2021; Millman et al., 2020; Whiteley et al., 2019). During phage infection, these enzymes are activated and produce cyclic oligonucleotides that bind to and activate a downstream effector protein. The activated effector proteins are proposed to induce premature cell death through various mechanisms, including membrane impairment (Cohen et al., 2019; Duncan-Lowey et al., 2021), DNA degradation (Fatma et al., 2021; Lau et al., 2020; Lowey et al., 2020), NAD^+^ depletion (Ko et al., 2022; Morehouse et al., 2022), amongst others. This anti-phage strategy was named <u>c</u>yclic-oligonucleotide-<u>b</u>ased <u>a</u>nti-phage <u>s</u>ignaling <u>s</u>ystem (CBASS).

As a countermeasure to CBASS immunity, phages encode anti-CBASS proteins. Acb1 degrades the cyclic nucleotide messengers to inhibit CBASS (Hobbs et al., 2022) and Acb2 is a cyclic dinucleotide (CDN) sponge (Huiting et al., 2023; Jenson et al., 2023). Interestingly, Acb2 also binds to a variety of other CDNs: 2’,3’-cGAMP and 3’,3’-cUU/UA/UG/AA with varying affinities. Structures of Acb2 from *P. aeruginosa* phage PaMx33 and from *E. coli* phage T4 in complex with 3’,3’-cGAMP showed that Acb2 forms an interlocked hexamer and binds to three cyclic dinucleotides, each with a binding pocket located in one Acb2 dimer within the hexamer (Huiting et al., 2023; Jenson et al., 2023). However, it remains unknown whether Acb2 binds to cyclic oligonucleotides that are utilized in Pycsar, CBASS and Type III CRISPR-Cas signaling systems (Molina et al., 2020; Tal et al., 2021; van Beljouw et al., 2022). Here, we find that Acb2 not only binds to and sequesters a broad spectrum of cyclic dinucleotides, but it also binds to cyclic trinucleotides 3’3’3’-cyclic AMP-AMP-AMP (cA3 hereafter) and 3’3’3’-cAAG (cAAG) with an order of magnitude higher affinity. CBASS systems commonly use these cyclic trinucleotide signals, as do some Type III CRISPR-Cas systems (Mayo-Muñoz et al., 2022). Structural characterization identified that one Acb2 hexamer binds two cyclic trinucleotides within binding pockets that are different from those binding cyclic dinucleotides. A co-structure of Acb2 bound to cA3 and 3’,3’-cGAMP at the same time is presented, as are mutants that independently disrupt the different binding sites. Phage-encoded Acb2 effectively protects phage when infecting cells simultaneously expressing cA3 and cGAMP CBASS systems. Together, this work identifies two distinct cyclic oligonucleotide binding sites on Acb2 and demonstrates that Acb2 is an effective, broad-spectrum inhibitor of CBASS and cGAS-STING signaling pathways.

## Results

### Acb2 sequesters diverse cyclic dinucleotides and is active in human cells

To understand the selectively of the newly identified Acb2 protein fold, we comprehensively tested an array of cyclic oligonucleotides that Acb2 may bind to. Previous work revealed that Acb2 binds to 3’,3’-cGAMP, 2’,3’-cGAMP and 3’,3’-cUU/UA/UG/AA, but not 3’,3’-cGG (Huiting et al., 2023) (Figure S1A-B). The study of Acb1 showed that, among cyclic dinucleotides, it does not cleave 3’,2’-cGAMP or 3’,3’-cGG/cUU (Hobbs et al., 2022).. In this current study, we first tested Acb2 binding of 3’,2’-cGAMP, which was recently identified as a signaling molecule for both CBASS and cGAS-like enzymes in eukaryotes (Fatma et al., 2021; Slavik et al., 2021). Interestingly, a native gel assay showed a significant shift of the Acb2 protein upon adding 3’,2’-cGAMP (Figure 1A), and isothermal calorimetry (ITC) experiments verified that Acb2 binds to 3’,2’-cGAMP with a *KD* of ~297.7 nM (Figures 1B and S2). Next, we solved the structure of Acb2 complexed with 3’,2’-cGAMP (2.33 Å), in which 3’,2’-cGAMP showed a similar binding mode as 3’,3’-cGAMP and c-di-AMP (Figures 1C-E and Table 1). Specifically, these dinucleotide molecules are bound by the N-terminal domains of the two interacting Acb2 protomers, and the π-π stacking from Y11 residue and salt bridges from K26 residue of both protomers further stabilizes this interaction (Figures 1D-E). The structure of Acb2 complexed with another cGAMP isomer, 2’,3’-cGAMP, solved at 2.24 Å resolution further confirmed this mode of binding (Figure 1E and Table 1). Since 3’,2’-cGAMP and 2’,3’-cGAMP are ligands used in eukaryotic cGAS-STING immunity, we also tested whether Acb2 can antagonize cGAS-STING signaling pathway in human cells. The results showed that upon expression of WT Acb2, interferon (IFN) signaling mediated by 2’,3’-cGAMP is significantly reduced while Y11A and K26A Acb2 mutants were less active (Figure 1F). Consistent with these data, native gel assays showed that Y11A and K26A mutations abrogated 2’,3’-cGAMP binding (Figure S1C). Taken together, our data demonstrates that Acb2 harbors a binding pocket that is well suited for many cyclic dinucleotides.

**Figure 1.**
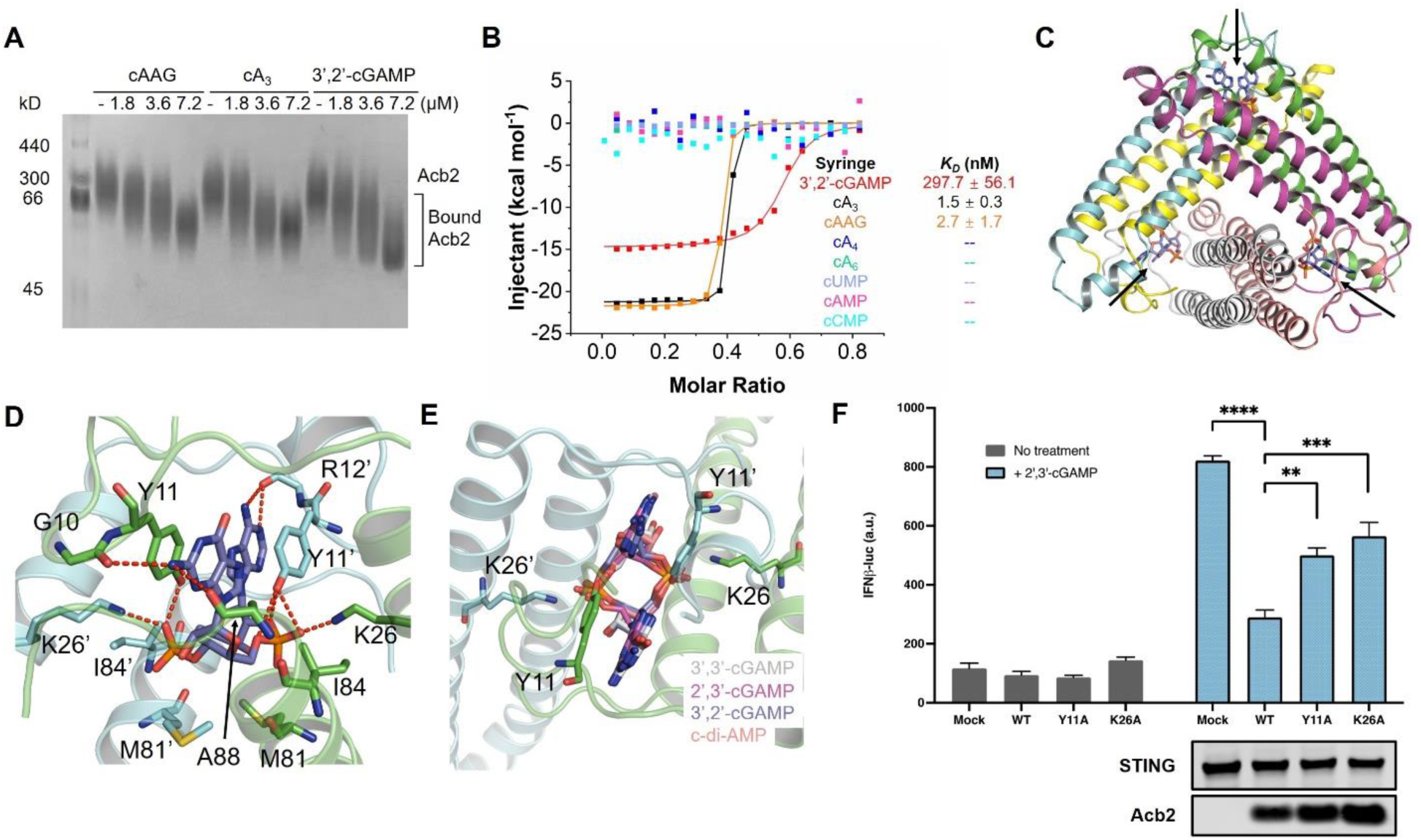
PaMx33 Acb2 also binds cyclic trinucleotides and 3’, 2’-cGAMP. (A) Native PAGE showed the binding of PaMx33-Acb2 to cyclic oligonucleotides. (B) ITC assays to test binding of cyclic nucleotides to PaMx33-Acb2. Representative binding curves and binding affinities are shown. The *KD* values are mean ± s.d. (n=3). Raw data for these curves are shown in Figure S2. (C) Overall structure of Acb2 complexed with 3’,2’-cGAMP, which are indicated by arrows. (D) Detailed binding between Acb2 and 3’,2’-cGAMP. Residues involved in 3’,2’-cGAMP binding are shown as sticks. Red dashed lines represent polar interactions. (E) Structural alignment among 3’,2’-cGAMP, 2’,3’-cGAMP, 3’,3’-cGAMP and c-di-AMP bound Acb2, highlighting the binding pocket of cyclic dinucleotides. (F) 293T-Dual cells were transfected with hSTING and Acb2 or its mutants, then treated with 2’,3’-cGAMP. STING activation was read as luciferase signal controlled by an interferon promoter.

**Table 1.**
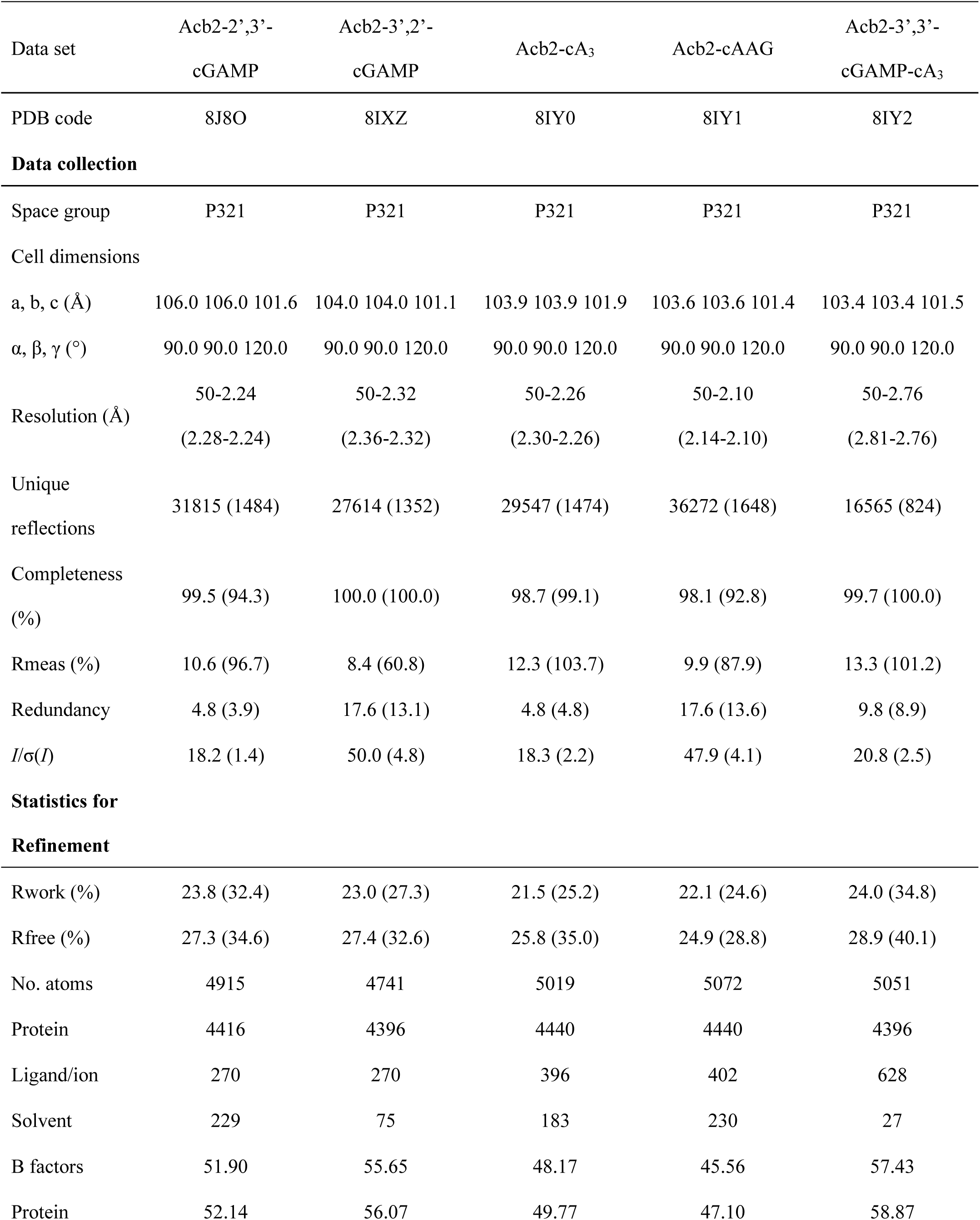
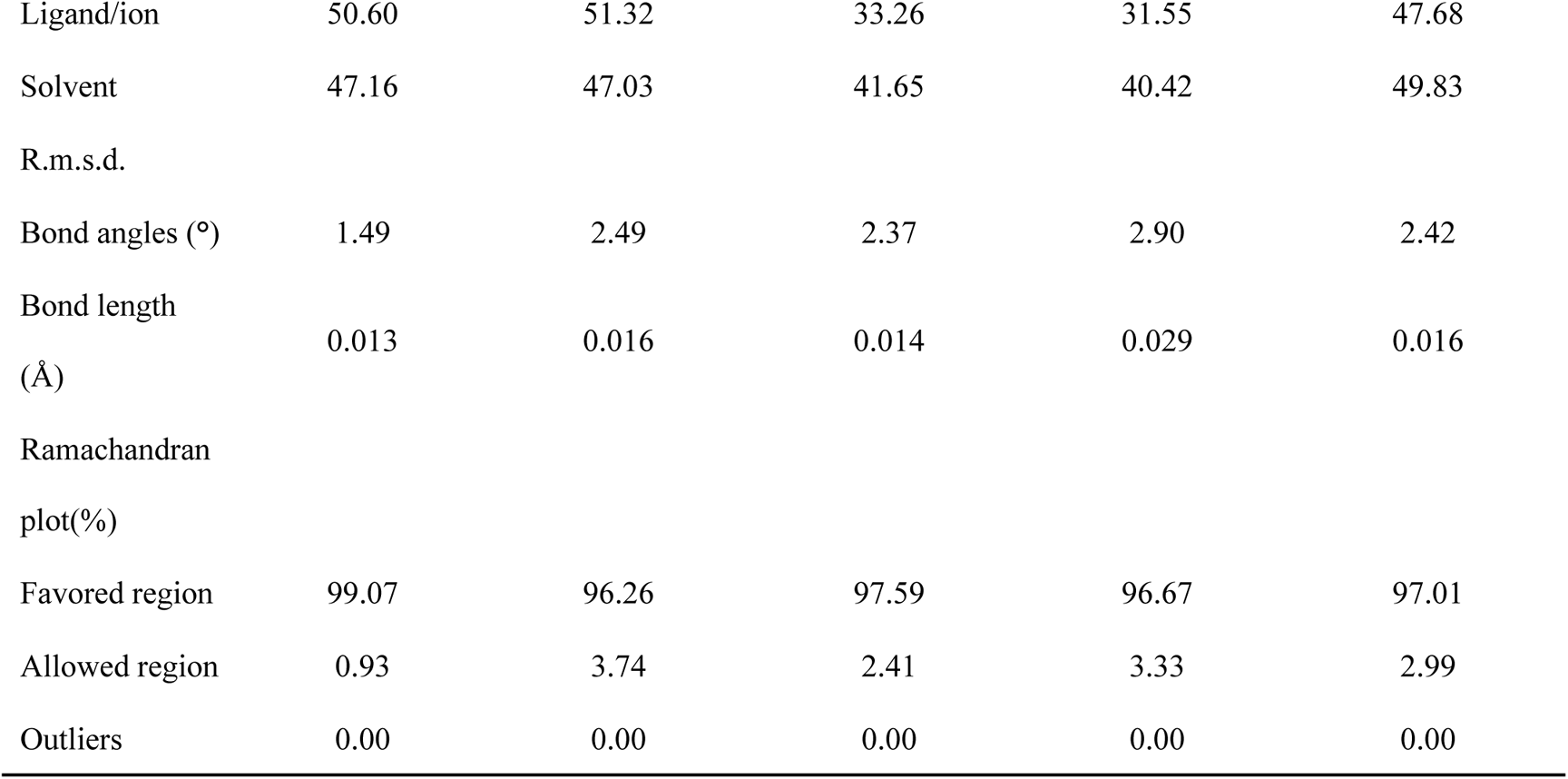
Data collection and refinement statistic.

| Data set | Acb2-2',3'-<br>cGAMP |  |  | Acb2-3',2'-<br>cGAMP |  |  | Acb2-cA<sub>3</sub> |  |  | Acb2-cAAG |  |  | Acb2-3',3'-<br>cGAMP-cA<sub>3</sub> |  |  |
| --- | --- | --- | --- | --- | --- | --- | --- | --- | --- | --- | --- | --- | --- | --- | --- |
| PDB code | 8J8O |  |  | 8IXZ |  |  | 8IY0 |  |  | 8IY1 |  |  | 8IY2 |  |  |
| Data collection |  |  |  |  |  |  |  |  |  |  |  |  |  |  |  |
| Space group | P321 |  |  | P321 |  |  | P321 |  |  | P321 |  |  | P321 |  |  |
| Cell dimensions |  |  |  |  |  |  |  |  |  |  |  |  |  |  |  |
| a, b, c (Å) | 106.0 | 106.0 | 101.6 | 104.0 | 104.0 | 101.1 | 103.9 | 103.9 | 101.9 | 103.6 | 103.6 | 101.4 | 103.4 | 103.4 | 101.5 |
| α, β, γ (°) | 90.0 | 90.0 | 120.0 | 90.0 | 90.0 | 120.0 | 90.0 | 90.0 | 120.0 | 90.0 | 90.0 | 120.0 | 90.0 | 90.0 | 120.0 |
| Resolution (Å) | 50-2.24 |  |  | 50-2.32 |  |  | 50-2.26 |  |  | 50-2.10 |  |  | 50-2.76 |  |  |
|  | (2.28-2.24) |  |  | (2.36-2.32) |  |  | (2.30-2.26) |  |  | (2.14-2.10) |  |  | (2.81-2.76) |  |  |
| Unique reflections | 31815 (1484) |  |  | 27614 (1352) |  |  | 29547 (1474) |  |  | 36272 (1648) |  |  | 16565 (824) |  |  |
| Completeness (%) | 99.5 (94.3) |  |  | 100.0 (100.0) |  |  | 98.7 (99.1) |  |  | 98.1 (92.8) |  |  | 99.7 (100.0) |  |  |
| Rmeas (%) | 10.6 (96.7) |  |  | 8.4 (60.8) |  |  | 12.3 (103.7) |  |  | 9.9 (87.9) |  |  | 13.3 (101.2) |  |  |
| Redundancy | 4.8 (3.9) |  |  | 17.6 (13.1) |  |  | 4.8 (4.8) |  |  | 17.6 (13.6) |  |  | 9.8 (8.9) |  |  |
| *I*/σ(*I*) | 18.2 (1.4) |  |  | 50.0 (4.8) |  |  | 18.3 (2.2) |  |  | 47.9 (4.1) |  |  | 20.8 (2.5) |  |  |
| Statistics for Refinement |  |  |  |  |  |  |  |  |  |  |  |  |  |  |  |
| Rwork (%) | 23.8 (32.4) |  |  | 23.0 (27.3) |  |  | 21.5 (25.2) |  |  | 22.1 (24.6) |  |  | 24.0 (34.8) |  |  |
| Rfree (%) | 27.3 (34.6) |  |  | 27.4 (32.6) |  |  | 25.8 (35.0) |  |  | 24.9 (28.8) |  |  | 28.9 (40.1) |  |  |
| No. atoms | 4915 |  |  | 4741 |  |  | 5019 |  |  | 5072 |  |  | 5051 |  |  |
| Protein | 4416 |  |  | 4396 |  |  | 4440 |  |  | 4440 |  |  | 4396 |  |  |
| Ligand/ion | 270 |  |  | 270 |  |  | 396 |  |  | 402 |  |  | 628 |  |  |
| Solvent | 229 |  |  | 75 |  |  | 183 |  |  | 230 |  |  | 27 |  |  |
| B factors | 51.90 |  |  | 55.65 |  |  | 48.17 |  |  | 45.56 |  |  | 57.43 |  |  |
| Protein | 52.14 |  |  | 56.07 |  |  | 49.77 |  |  | 47.10 |  |  | 58.87 |  |  |

|  |  |  |  |  |  |
| --- | --- | --- | --- | --- | --- |
| Ligand/ion | 50.60 | 51.32 | 33.26 | 31.55 | 47.68 |
| Solvent | 47.16 | 47.03 | 41.65 | 40.42 | 49.83 |
| R.m.s.d. |  |  |  |  |  |
| Bond angles (°) | 1.49 | 2.49 | 2.37 | 2.90 | 2.42 |
| Bond length<br>(Å) | 0.013 | 0.016 | 0.014 | 0.029 | 0.016 |
| Ramachandran<br>plot(%) |  |  |  |  |  |
| Favored region | 99.07 | 96.26 | 97.59 | 96.67 | 97.01 |
| Allowed region | 0.93 | 3.74 | 2.41 | 3.33 | 2.99 |
| Outliers | 0.00 | 0.00 | 0.00 | 0.00 | 0.00 |

### Acb2 sequesters cyclic trinucleotides with higher affinity than cyclic dinucleotides

Based on the binding pocket of Acb2, we hypothesized that Acb2 may not bind cyclic mononucleotides or oligonucleotides, such as cA3, cA4 or cA6, due to potential steric clash caused by the nucleotides. Of note, cA3 and cAAG are major products of the CD-NTase enzymes involved in CBASS whereas cA4 is only a minor product of a single CD-NTase (Lowey et al., 2020). cA6 has not been identified as a product of any known CD-NTases. However, all three cyclic oligoadenylates are known products involved in Type III CRISPR-Cas anti-phage immunity (Athukoralage and White, 2022). A native gel assay showed that the Acb2 protein does not shift upon adding cA4 or cA6 molecules (Figures S1A-B). Furthermore, both native gel and ITC assays showed that Acb2 does not bind to cUMP, cCMP or cAMP (Figures S1-2 and 1B). However, the native gel assay revealed a significant shift of the Acb2 protein upon adding cA3 or cAAG. ITC experiments revealed that Acb2 binds to cA3 and cAAG with a *KD* of ~1.5 and ~2.7 nM (Figures 1B and S2), respectively, which is more than an order of magnitude stronger than Acb2 binding to 3’,3’-cGAMP (*KD* of ~87 nM) (Huiting et al., 2023). To determine whether Acb2 sequesters or cleaves the cyclic trinucleotide molecules, high-performance liquid chromatography (HPLC) revealed that incubating Acb2 with cA3 depletes any detectable molecules, and following proteolysis of Acb2, cA3 is released back into the buffer unmodified (Figure 2A). Collectively, these results demonstrate that Acb2 binds to and sequesters cyclic trinucleotides commonly used in CBASS immunity with a significantly higher affinity than cyclic dinucleotides.

**Figure 2.**
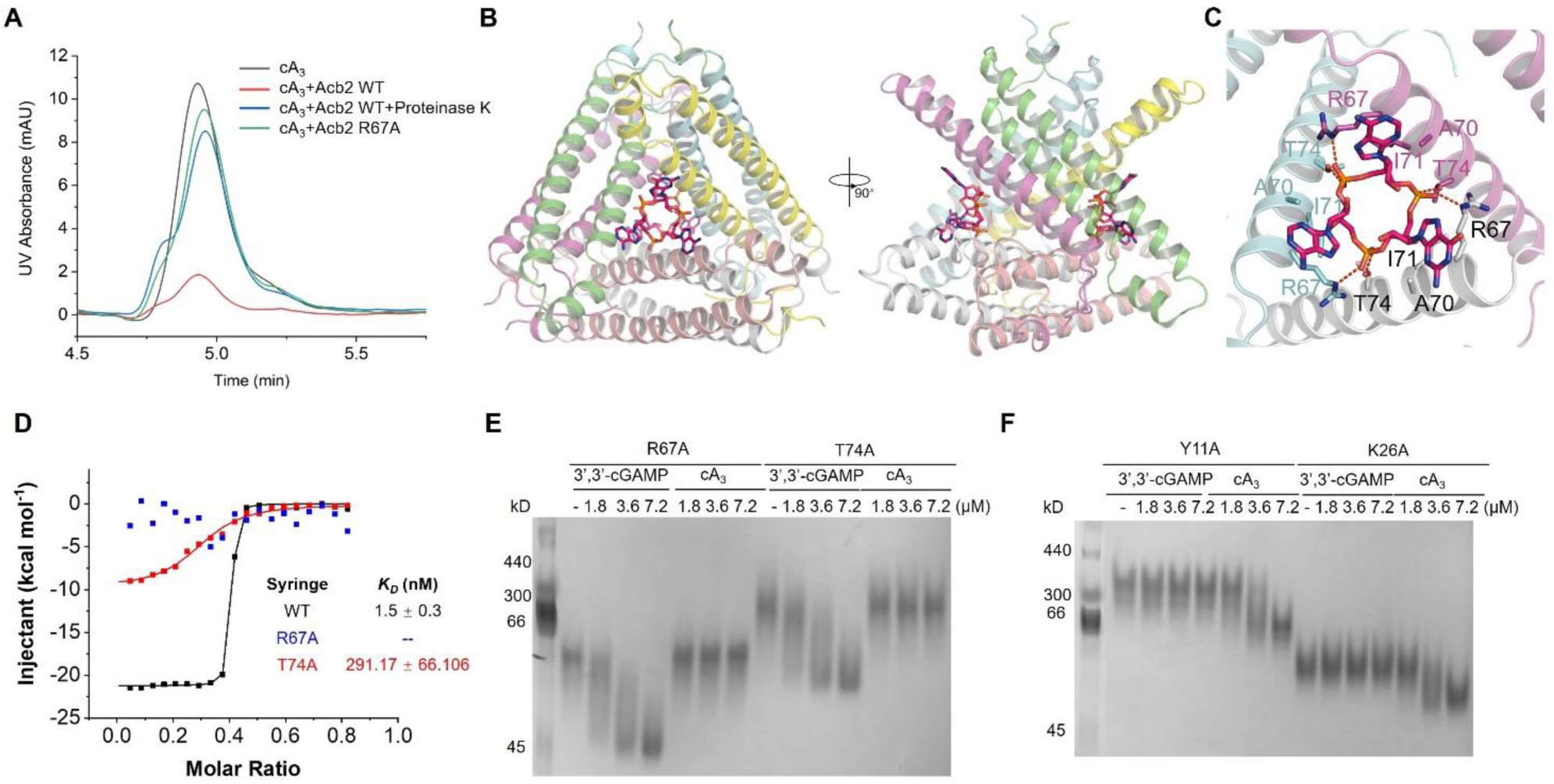
Acb2 binds to cyclic trinucleotides with binding sites different from those of cyclic dinucleotides. (A) The ability of PaMx33-Acb2 to bind and release cA3 when treated with proteinase K was analyzed by HPLC. cA3 standard was used as a control. The remaining cA3 after incubation with PaMx33-Acb2 was tested. (B) Overall structure of Acb2 complexed with cAAG, which are shown as sticks. Two views are shown. (C) Detailed binding between Acb2 and cAAG. Residues involved in cAAG binding are shown as sticks. Red dashed lines represent polar interactions. (D) ITC assays to test the binding of cA3 to PaMx33-Acb2 mutants. Representative binding curves and binding affinities are shown. The *KD* values are mean ± s.d (n = 3). Raw data for these curves are shown in Figure S2. (E-F) Native PAGE showed the binding of PaMx33 Acb2 mutants to cyclic oligonucleotides.

### Acb2 binds cyclic trinucleotides and dinucleotides with different binding sites

The binding of cyclic trinucleotides was unexpected because the Acb2 binding pocket appears well suited for only cyclic dinucleotides. To understand how Acb2 interacts with cyclic trinucleotides, we determined the crystal structures of Acb2 in complex with cA3 (2.26 Å) or cAAG (2.10 Å) (Table 1). Surprisingly, the structures showed that one Acb2 hexamer binds two cyclic trinucleotides with two distinct binding pockets that are far from the three pockets for the cyclic dinucleotide binding (Figures 2B and S3A-B). Each binding site of the cyclic trinucleotides is formed by three Acb2 protomers in a three-fold symmetry and the two binding sites are opposite each other (Figure 2B). In turn, each of the two protomers that together bind a cyclic dinucleotide is involved in binding to one out of the two cyclic trinucleotides, respectively (Figure S3A). Correspondingly, each of the three protomers that together bind a cyclic trinucleotide is involved in binding to one out of the three cyclic dinucleotides, respectively (Figure S3B).

The cyclic trinucleotide is bound mainly through its three phosphate groups, each of which is coordinated by R67 of one protomer and T74 of another protomer through hydrogen bonds (Figure 2C). Moreover, the cyclic trinucleotide is also stabilized by hydrophobic interactions from R67, A70 an I71 from each of the three protomers (Figure 2C). Consistent with this analysis, the Acb2 T74A mutant displayed a significantly decreased binding affinity to cA3 (*KD* of ~291 nM), and the Acb2 R67A mutant abolished Acb2 binding of cA3 *in vitro* (Figures 2D and S2). To confirm that the binding sites of the cyclic trinucleotides and dinucleotides in Acb2 are independent of each other, we tested the binding of 3’,3’-cGAMP with the T74A or R67A Acb2 mutant proteins. A native gel assay showed similar shifts of the two Acb2 mutants as WT Acb2 upon adding 3’,3’-cGAMP (Figure 2E), suggesting that the binding to 3’,3’-cGAMP is not affected by the two mutations. In turn, we tested the binding of cA3 with Y11A and K26A Acb2 mutants, which lose their binding to 3’,3’-cGAMP (Huiting et al., 2023). The native gel results showed a significant shift of Y11A and K26A mutant proteins upon adding cA3 (Figure 2F). Taken together, these data collectively show that one Acb2 hexamer binds two cyclic trinucleotides through two pockets independent of those that bind cyclic dinucleotides.

Structural alignment between apo Acb2 and its complexes with cyclic trinucleotides showed that the binding of cyclic trinucleotides does not induce a conformational change of Acb2, with a root mean square deviation (RMSD) of 0.224 and 0.261 Å (Cα atoms) for Acb2-cA3 and Acb2-cAAG compared to the apo Acb2, respectively (Figure S3C). Therefore, we co-crystallized Acb2 with both cA3 and 3’,3’-cGAMP and then solved its crystal structure at a resolution of 2.76 Å (Table 1). The structure clearly showed that Acb2 binds to two cA3 and three 3’,3’-cGAMP molecules simultaneously (Figures 3A-C). Structural alignment between Acb2-cA3-3’,3’-cGAMP and apo Acb2 also showed little conformational changes with an RMSD of 0.298 Å for Cα atoms (Figure S3D).

**Figure 3.**
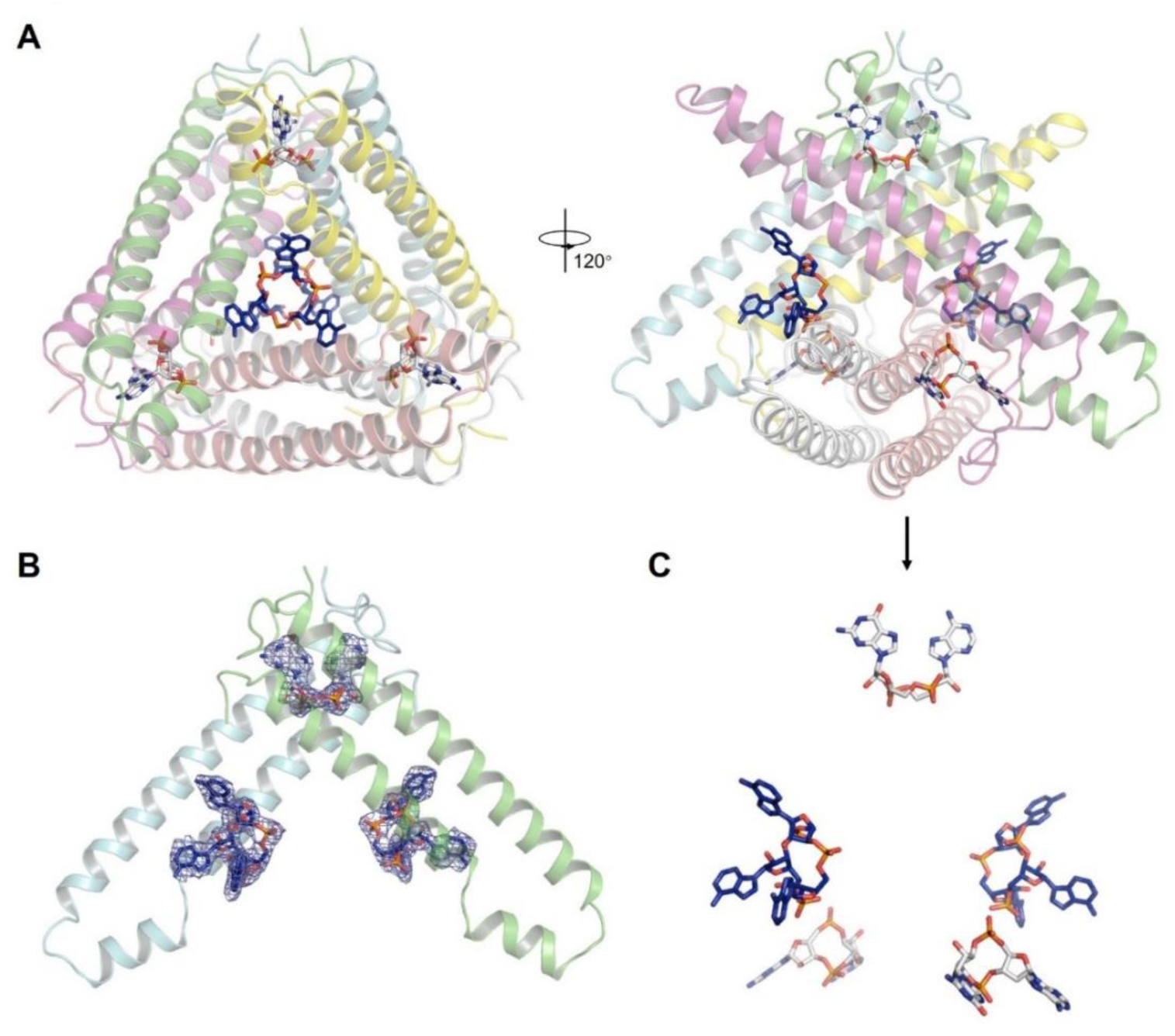
Acb2 binds to cyclic trinucleotides and dinucleotides simultaneously. (A) Overall structure of Acb2 complexed with cA3 and 3’,3’-cGAMP. cA3 and 3’,3’-cGAMP are shown as blue and light gray sticks. Two views are shown. (B) 2Fo-Fc electron density of cA3 and 3’,3’-cGAMP within an Acb2 dimer contoured at 1 σ. (C) Distribution of the small molecules within the Acb2 hexamer. The nucleotides are shown as they are in the right panel of (A).

### Acb2 binds to cA_3_ with a novel fold

Dali search did not return entries of experimentally determined proteins with the same fold as Acb2, nor did Foldseek searches of computationally predicted proteins (van Kempen et al., 2023), suggesting that both the cyclic dinucleotide and trinucleotide-binding folds are novel. Several experimentally determined proteins have been reported to bind cyclic trinucleotides, including the CBASS effector proteins NucC (Lau et al., 2020) and Cap4 (Lowey et al., 2020) that directly bind cA3, as well as the human CDN sensor RECON that directly binds cAAG (Whiteley et al., 2019). Compared to the cA3 binding pocket in Acb2, those in NucC, Cap4, and RECON are significantly different. In NucC, one cA3 molecule is bound in a three-fold symmetric allosteric pocket at the ‘‘bottom’’ of the protein trimer, mainly formed by an extended hairpin loop from each protomer. Additionally, each adenine base is stabilized by hydrogen bonds and π stacking interactions in NucC (Figure S4A, PDB code: 6Q1H). In Cap4, cA3 is bound within its SAVED (SMODS-associated) domain, which is a fusion of two CARF (CRISPR-associated Rossman fold) domains derived from Type III CRISPR-Cas system (Figure S4B, PDB code: 6WAN). RECON adopts a TIM barrel fold with eight parallel β strands surrounded by eight crossover α-helixes and cAAG is bound in a deep crevice at the top of the β barrel (Figure S4C, PDB code: 6M7K). Moreover, the conformation of cA3 within Acb2 is also different from those within NucC, Cap4, and RECON complex structures (Figure S4D). Specifically, cA3 in both NucC and Cap4 are almost in an overall planar conformation, and two adenine bases of cAAG within RECON are nearly in the same plane as the phosphodiester ring and the third guanine base is extended out. However, each base of cA3 forms a ~46.8 degree angle with the phosphate plane in Acb2. Together, the structure of Acb2 complexed with cA3 reveals a novel cyclic trinucleotide-binding fold.

### Cyclic nucleotide binding spectra are different among Acb2 homologs

Compared to the highly conserved Y11 and K26 residues of Acb2 (essential for 3’,3’-cGAMP binding) found in the *Pseudomonas aeruginosa* phage PaMx33 protein, the R67 and T74 residues (essential for cA3 binding) are less conserved and vary in some Acb2 homologs (Figure 4A) (Huiting et al., 2023). Therefore, we assessed the binding spectrum of other Acb2 homologs to determine whether the homologs bind to the same spectrum of cyclic oligonucleotides. We chose Acb2 homologs from *P. aeruginosa* phage JBD67 (44.4% a.a. identity), in which both R67 and T74 residues are conserved, *Serratia* phage CHI14 (23.5% a.a. identity) and *Escherichia* phage T4 (24.2% a.a. identity), in which only R67 (Serratia phage) or T74 (Escherichia phage) residue is conserved. ITC analyses showed that JBD67-Acb2 directly binds to 3’,3’-cGAMP with a *KD* of ~99 nM and cA3 with a *KD* of ~3.5 nM (Figure 4B and S5A-B), both of which are comparable to those of PaMx33-Acb2. Native gel assays also suggest that JBD67-Acb2 binds to the same spectrum of cyclic nucleotides as PaMx33-Acb2 (Figure S6A). ITC analyses showed that T4-Acb2 directly binds to 3’,3’-cGAMP with a *KD* of ~84.4 nM, consistent with previous work (Jenson et al., 2023), but does not bind to cA3 (Figure 4C and S5C-D). Native gel assays also suggest that T4-Acb2 binds the same spectrum of cyclic dinucleotides as PaMx33-Acb2, but not to the cyclic trinucleotides cA3 and cAAG (Figure S6B). Next, we mutated D61 of T4-Acb2 to Arginine to see whether it can endow T4-Acb2 with the binding activity of cA3 because T4-Acb2 already has T68 residue in the place of T74 of PaMx33-Acb2 that is essential for cA3 binding. However, based on ITC assays, we observed that the D61R mutant of T4-Acb2 was still unable to bind cA3 (Figures 4C and S5E). Interestingly, structural alignment between T4-Acb2 (Jenson et al., 2023) and PaMx33-Acb2 complexed with cA3 showed that the helix lining the cA3 binding pocket of PaMx33-Acb2 has a kink at E64, which enlarges the pocket to accommodate the base groups of cA3 (Figure 4D). However, the corresponding helix of T4-Acb2 does not kink here, so the Y37, A57, L60, D61 and T64 residues of T4-Acb2 may undergo steric clashing and prevent cA3 binding (Figure 4D). More importantly, the relative angles among the three helices lining the binding pocket are also different between PaMx33-Acb2 and T4-Acb2, resulting in a smaller binding pocket in T4-Acb2 (Figure 4E). Together, these observations may explain the inability of T4-Acb2 binding to cyclic trinucleotides. Lastly, ITC analyses showed that CHI14-Acb2 directly binds to 3’,3’-cGAMP with a *KD* of ~62.4 nM, but does not bind to cA3 (Figure 4F). Of note, native gel assays showed almost no shift of the CHI14-Acb2 protein upon adding any cyclic oligonucleotides, including 3’,3’-cGAMP (Figure S6C), suggesting that native gel assay is not suitable for studying the binding spectrum of CHI14-Acb2. Therefore, we studied the binding of all the cyclic oligonucleotides to CHI14-Acb2 through ITC analyses, which showed that CHI14-Acb2 exhibits the same binding spectrum as T4-Acb2 (Figures 4F-G and S7). In summary, Acb2 homologs bind to many cyclic trinucleotides and dinucleotides used in cGAS-based immunity with certain homologs being more broad spectrum than enzyme Acb1 (Figure 4H).

**Figure 4.**
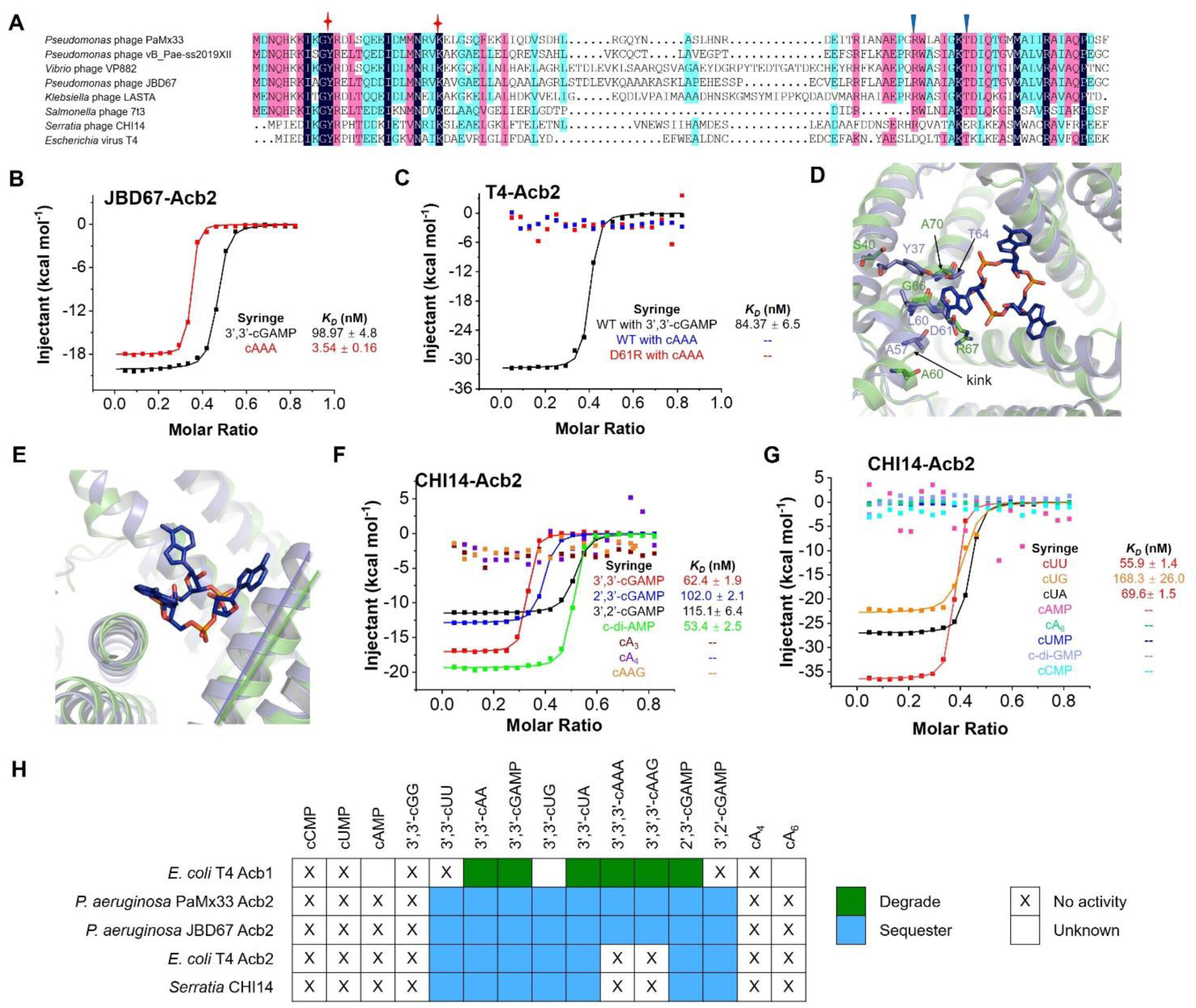
The binding spectra are different among Acb2 homologs. (A) Sequence alignment among Acb2 homologs. Residues with 100 % identity, over 75 % identity and over 50 % identity are shaded in dark blue, pink and cyan, respectively. Residues involved in binding of cyclic dinucleotides and trinucleotides are marked with stars and triangles, respectively. (B) ITC assays to test binding of cyclic oligonucleotides to JBD67-Acb2. Representative binding curves and binding affinities are shown. The *KD* values are mean ± s.d. (n=3). Raw data for these curves are shown in Figure S5. (C) ITC assays to test binding of cyclic oligonucleotides to T4-Acb2. Representative binding curves and binding affinities are shown. The *KD* values are mean ± s.d. (n=3). Raw data for these curves are shown in Figure S5. (D) Structural alignment between PaMx33-Acb2 and T4-Acb2 at one monomer. Residues with potential steric clash with cA3 in T4-Acb2 and the corresponding residues in PaMx33-Acb2 are shown in sticks. (E) The same alignment shown in (D), highlighting the different relative angels formed by the three helices lining the binding pocket of cA3. (F-G) ITC assays to test binding of cyclic nucleotides to CHI14-Acb2. Representative binding curves and binding affinities are shown. The *KD* values are mean ± s.d. (n=3). Raw data for these curves are shown in Figure S6. (H) Summary of the binding results of Acb2 homologs and Acb1 (Hobbs et al., 2022).

### Acb2 antagonizes Type III-C CBASS immunity

Since Acb2 displays high affinity binding to cyclic trinucleotides, we tested whether phage-encoded Acb2 can inhibit Type III-C CBASS immunity that uses a cA3 signaling molecule to activate endonuclease (NucC) effector protein. NucC is a cyclic nucleotide-activated effector in both CBASS and Type III CRISPR-Cas systems, where it non-specifically degrades DNA and prematurely kill the host cell before phage can effectively replicate (Lau et al., 2020; Mayo-Muñoz et al., 2022; Ye et al., 2020). First, we set up an *in vitro* NucC activity assay using purified NucC from the *P. aeruginosa* strain ATCC 27853. While cA3 activates the DNA cleavage activity of NucC, adding WT Acb2 into this system significantly decreased NucC activity (Figure 5A). Moreover, following proteolysis of WT Acb2, the released cA3 molecule again activated the NucC activity (Figure 5A; last two lanes). The R67A and T74A Acb2 mutant proteins, which lost or exhibited decreased cA3 binding, displayed minimal inhibition of NucC activity. However, the Y11A and K26A Acb2 mutants, whose cyclic dinucleotide binding pockets are disrupted, inhibited NucC activity similarly to WT Acb2 (Figure 5A). These results demonstrate that Acb2 antagonizes Type III-C CBASS immunity *in vitro* through sequestering the cA3 molecule.

**Figure 5.**
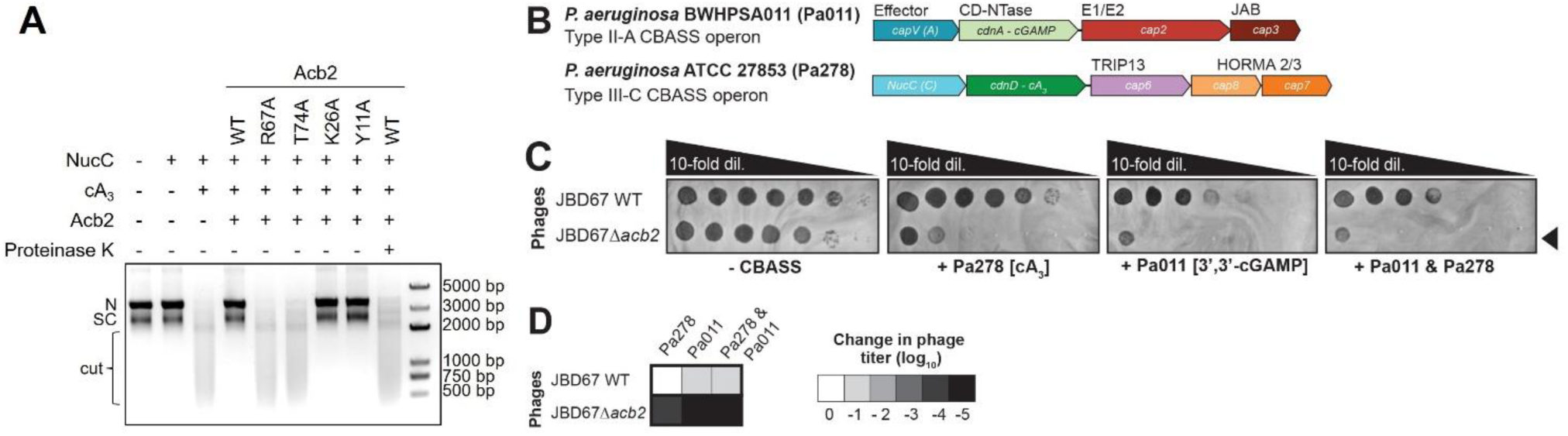
Acb2 antagonizes tri- and di-nucleotide based CBASS immunity. (A) Effect of Acb2 or its mutants on cA3-activated NucC effector protein function. After treatment with proteinase K, the released cA3 also showed the ability to activate the nuclease activity of NucC. The concentration of NucC, cA3, Acb2 and proteinase K is 10 nM, 5 nM, 50 nM and 1 µM, respectively. N denotes nicked plasmid, SC denotes closed-circular supercoiled plasmid, and cut denotes fully digested DNA. (B) *Pseudomonas aeruginosa* BWHPSA011 (Pa011) Type II-A CBASS and ATCC 27853 (Pa278) Type III-C CBASS operons. (C) Plaque assays with JBD67 WT and Δ*acb2* phages spotted in 10-fold serial dilutions on a lawn of PAO1 strain harboring an empty vector (E.V.) plasmid (-CBASS), PAO1 harboring a plasmid with the Pa278 CBASS operon (+ Pa278), PAO1 with a chromosomally integrated Pa011 CBASS operon (+ Pa011), and the aforementioned strain harboring a plasmid with the Pa278 CBASS operon (+ Pa011 & Pa278). Basal expression of both operons was sufficient to induce phage targeting (i.e. induction was not necessary). (D) Heat map representing the order of magnitude change in phage titer, where phage titer is quantified by comparing the number of spots (with plaques, or clearings if plaques were not visible) on the CBASS-expressing strains divided by the E.V.-expressing strain (n=3).

To determine whether phage-encoded Acb2 can inhibit this same cA3-based CBASS system *in vivo,* we performed phage infection assays with the *P. aeruginosa* ATCC 27853 strain harboring the Type III-C CBASS operon (Figure 5B). However, phages naturally expressing *acb2* were unable to replicate on the native *P. aeruginosa* strain, which is likely due to the presence of other anti-phage immune systems (Huiting et al. 2023), prophages, or receptor incompatibility. Next, we established a heterologous host expressing the *P. aeruginosa* ATCC 27853 (Pa278) Type III-C CBASS operon in a phage sensitive strain that naturally lacks CBASS (PAO1). We also used a *P. aeruginosa* strain with a chromosomally integrated BWHPSA011 (Pa011) Type II-A CBASS operon (3’,3’-cGAMP-producing), which Acb2 has been shown to antagonize (Huiting et al. 2023). In the presence of Pa278 Type III-C CBASS, WT JBD67 phage encoding Acb2 was unaffected, but the *acb2* mutant phage (JBD67Δ*acb2*) was reduced 4 orders of magnitude (Figures 5C-D), demonstrating potent Acb2 activity *in vivo* against a trinucleotide system. In the presence of the Pa011 Type II-A CBASS system, JBD67 WT phage titer was reduced by ~1 order of magnitude, consistent with Acb2 being partially overwhelmed, as shown previously (Huiting et al. 2023). This finding supports a hypothesis from Jenson et al., 2023 that proposes ubiquitin-like conjugation of the Type II-A CBASS CD-NTase enhances cGAMP production to overwhelm the CDN sponge (Jenson et al., 2023). By contrast, the titer of the *acb2* mutant phage was reduced 5 orders of magnitude (Figures 5C-D), which confirms the potent, yet incomplete, inhibition of Type II-A CBASS. Co-expression of Pa278 Type III-C CBASS (cA3 producing) and Pa011 Type II-A CBASS (3’,3’-cGAMP producing) targets the WT JBD67 phage to the same degree as Pa011 Type II-A CBASS alone (Figures 5C-D), demonstrating the versatility of Acb2 when simultaneously faced with two distinct CBASS systems *in vivo*.

## Discussion

Following phage infection, CBASS immunity functions via the activation of a cGAS-like enzyme to catalyze the synthesis of a cyclic oligonucleotide signaling molecule. To date, two phage proteins have been discovered to antagonize the CBASS immunity: Acb1 and Acb2. Acb1 uses an inhibitory mechanism common to the eukaryotic cGAS-STING signaling system (Eaglesham and Kranzusch, 2020), that is, enzymatically cleaving and depleting an array of cyclic dinucleotides and trinucleotides (Hobbs et al., 2022). In contrast, we, alongside another independent group, reported that Acb2 acts as a “sponge” and sequesters 3’,3’-cGAMP (Huiting et al., 2023; Jenson et al., 2023) as well as a variety of other CBASS cyclic dinucleotide signaling molecules (Huiting et al., 2023). A sponging mechanism was also reported for inhibitors of the anti-phage system Thoeris, including Tad1 (Leavitt et al., 2022) and Tad2 (Yirmiya et al., 2023). Here, we extend the dinucleotide binding spectrum to include 3’,2’-cGAMP, a signaling molecule not cleaved by Acb1, but recently implicated in both CBASS and cGAS like signaling in eukaryotes (Cai et al., 2023; Fatma et al., 2021; Slavik et al., 2021). The ability of Acb2 to function in human cells reinforces the flexibility of the “sponging” mechanism (i.e. no need to bind to CBASS proteins or host machinery), and the remarkable trans-kingdom conservation of this cyclic-oligonucleotide-based immune system. These data also open up the possibility that pathogenic bacteria encoding Acb1, Acb2, or other undiscovered cGAMP interactors (i.e. on prophages) could use them to dampen the human immune response during intracellular infection. Indeed, cross-kingdom signaling has been previously reported with bacterial c-di-GMP (Burdette et al., 2011) and c-di-AMP produced by *Listeria monocytogenes* (Woodward et al., 2010), which both serve as ligands for human STING. Taken together, these observations suggests that the interface between the bacterial signals, cyclic oligonucleotide inhibitors, and cGAS-STING immunity has yet to be fully explored.

Acb2 binds to and sequesters cyclic trinucleotides with a magnitude higher binding affinity than to dinucleotides. Based on this finding, we further observed that Acb2 binds to a wider spectrum of cyclic oligonucleotides compared to the enzyme Acb1 including 3’,3’-cUU, 3’,3’-cUG, and 3’,2’-cGAMP (Figure 4H). Moreover, Acb2 uses distinct, independent binding pockets to sequester cyclic dinucleotides and trinucleotides whereas Acb1 uses the same active site to degrade cyclic oligonucleotides. In some cases, however, we observed that Acb2 homologs cannot bind to cyclic trinucleotides. Although it is unclear if there is a cost to the trinucleotide binding site that would lead to its loss, in the case of phage T4, its genome encodes both Acb1 and Acb2 and therefore suggests that this phage is broadly evasive of most CBASS systems. To our knowledge, Acb2 represents the first protein that can simultaneously bind two types of cyclic oligonucleotides with different binding pockets. Altogether, our work highlights Acb2 as an effective and potent inhibitor of nearly all CBASS types, with great potential to fortify phage therapeutics by enabling evasion of most cGAS-based anti-phage systems in bacteria.

## Acknowledgments

We thank the staff at beamlines BL02U1 and BL19U1 of the Shanghai Synchrotron Radiation Facility for their assistance with data collection. We thank the Tsinghua University Branch of China National Center for Protein Sciences Beijing and Dr. Shilong Fan for providing facility support for X-ray diffraction of the crystal samples. We thank Drs. Yuanyuan Chen, Zhenwei Yang, Bingxue Zhou at the Institute of Biophysics, Chinese Academy of Sciences for technical help with ITC experiments. Y. F. is supported by National key research and development program of China (2022YFC3401500 and 2022YFC2104800), the National Natural Science Foundation of China (32171274) and Beijing Nova Program. E.H. is supported by the National Science Foundation Graduate Research Fellowship Program [Grant No. 2038436]. Any opinions, findings, and conclusions or recommendations expressed in this material are those of the authors and do not necessarily reflect the views of the National Science Foundation. J.B.-D. is supported by the National Institutes of Health [R21AI168811, R01GM127489], the Vallee Foundation, and the Searle Scholarship.

## Author contributions

Y.F. and J.B.-D. conceived and supervised the project and designed experiments. Xueli Cao, D. L., Y. W. and L. G. purified the proteins, grew and optimized the crystals, collected the diffraction data, and performed *in vitro* activity analysis and binding assays. Y.X. solved the crystal structures. E.H. performed all *in vivo* phage experiments. J.R. performed the HPLC analysis. Xujun Cao performed *in vivo* human cell experiments under supervision of L.L.. Y.F. wrote the original manuscript. J.B.-D., Y.F., and E.H. revised the manuscript.

## STAR Methods

### Key Resources Table

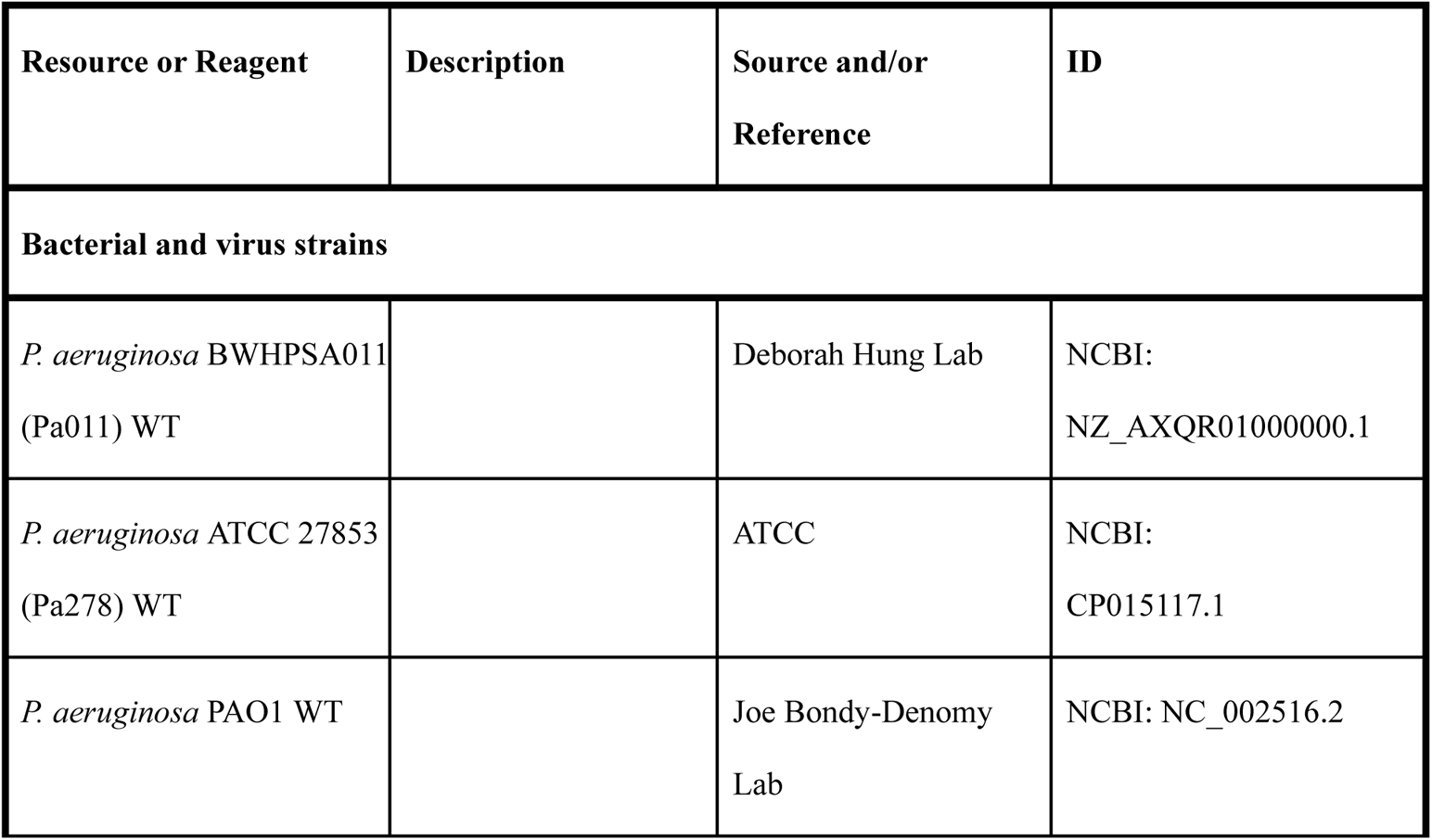
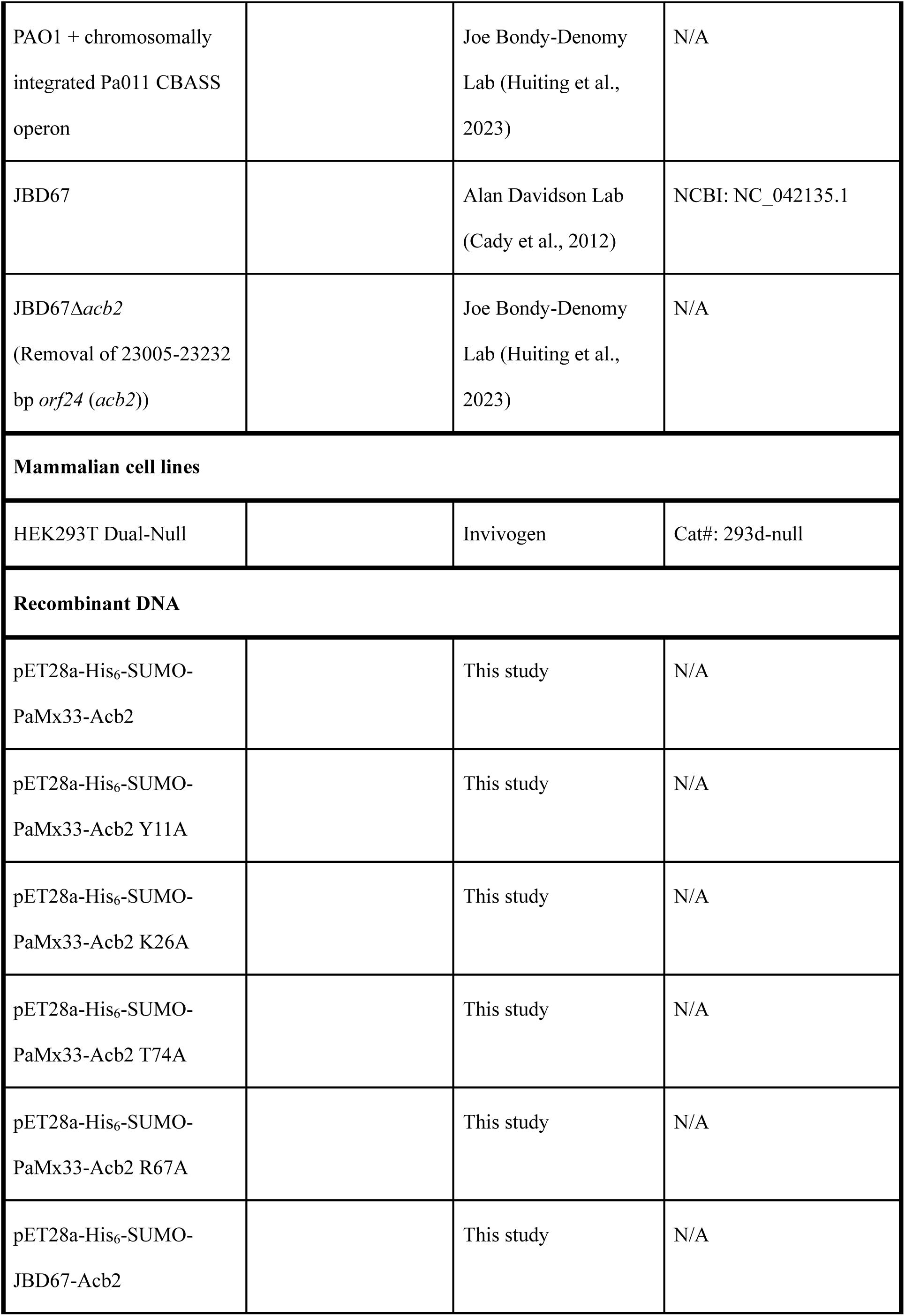
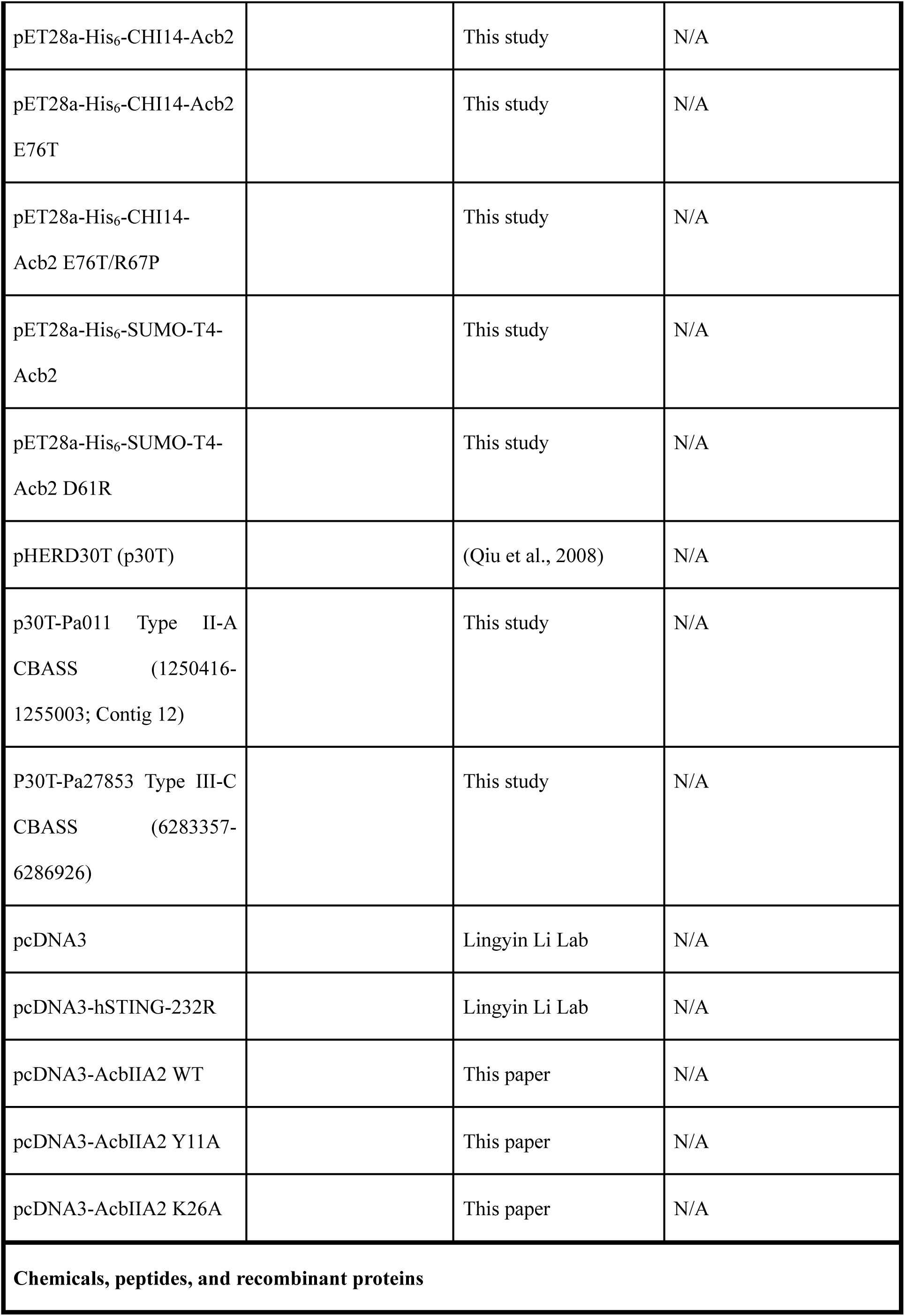
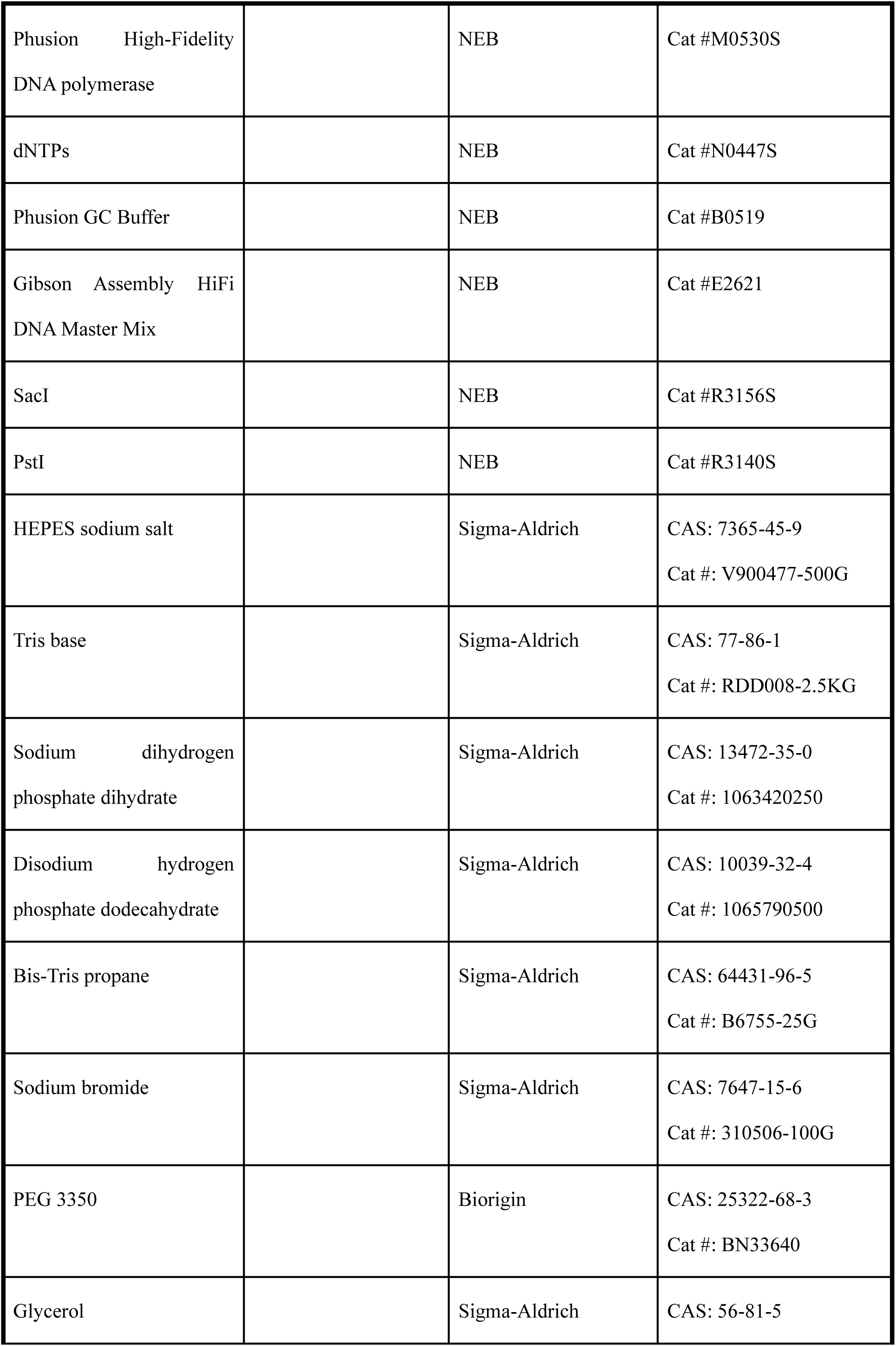
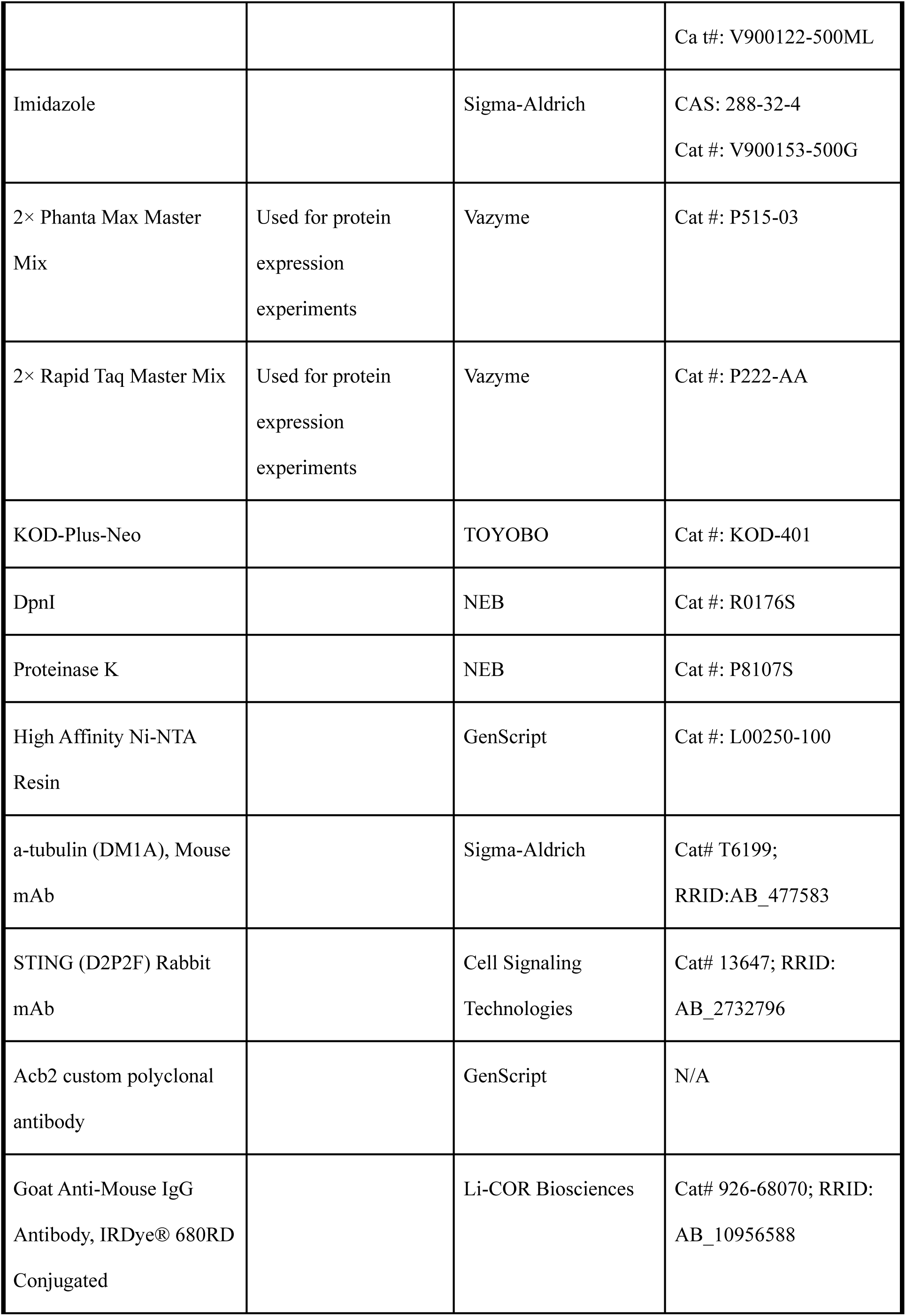
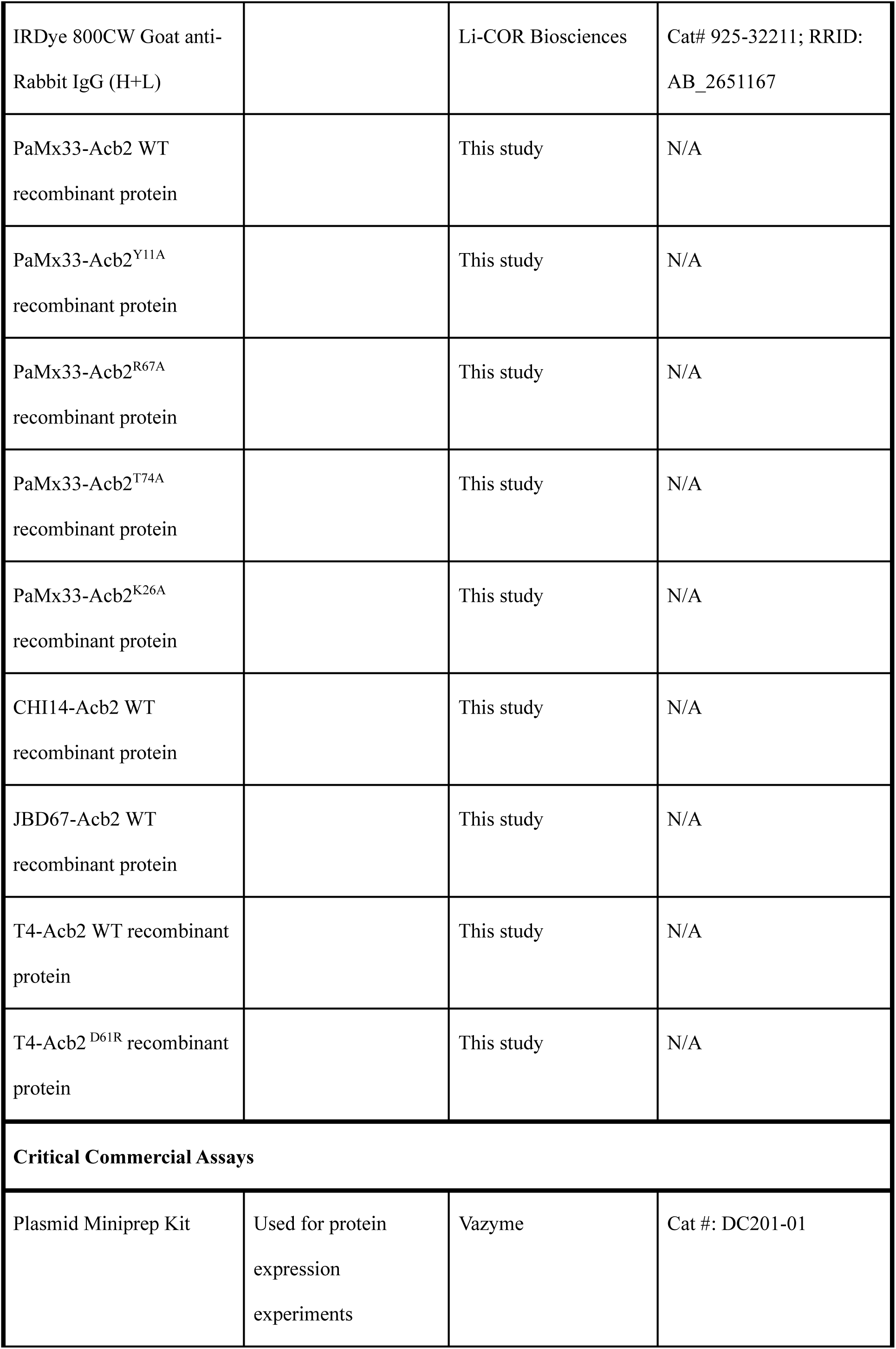
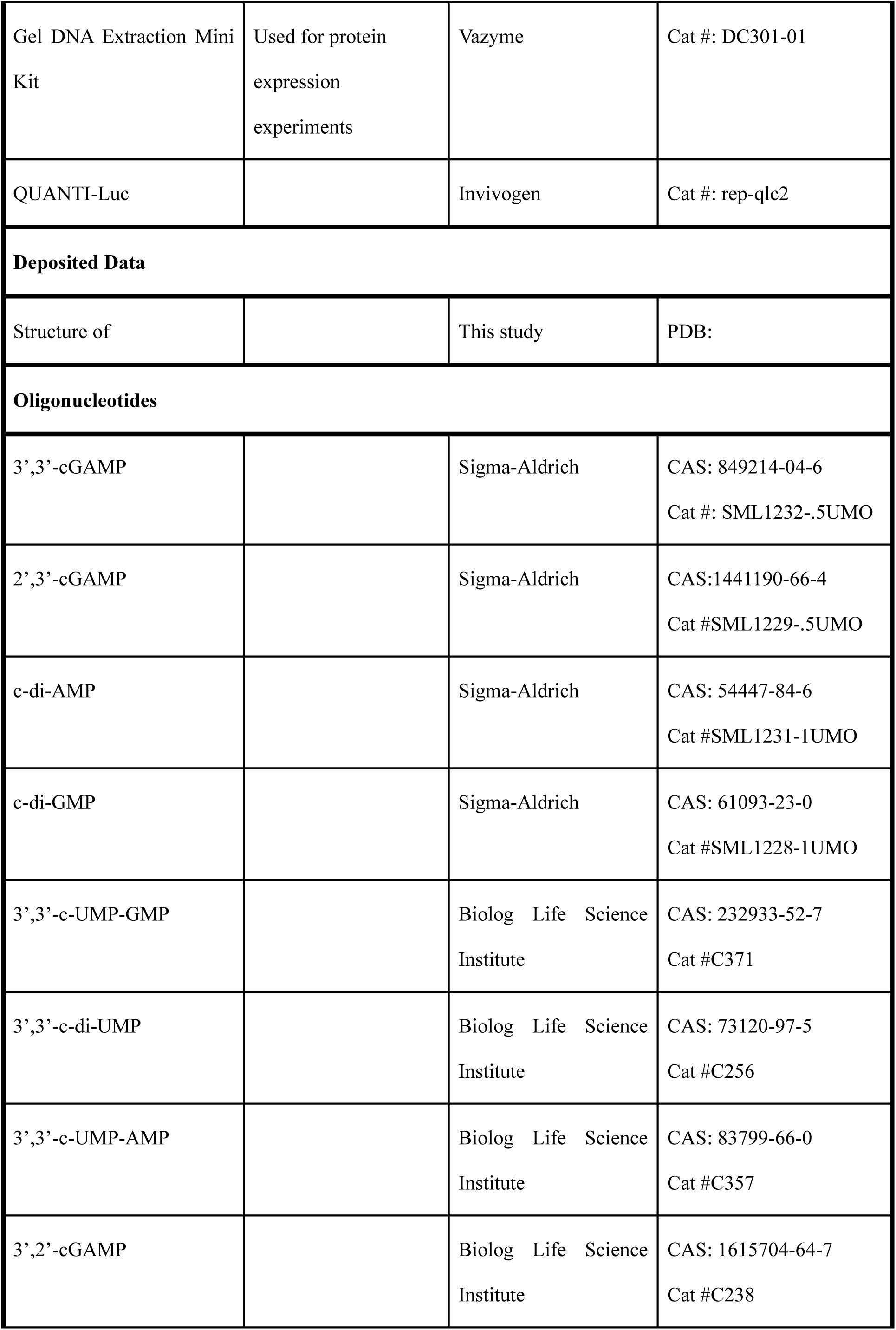
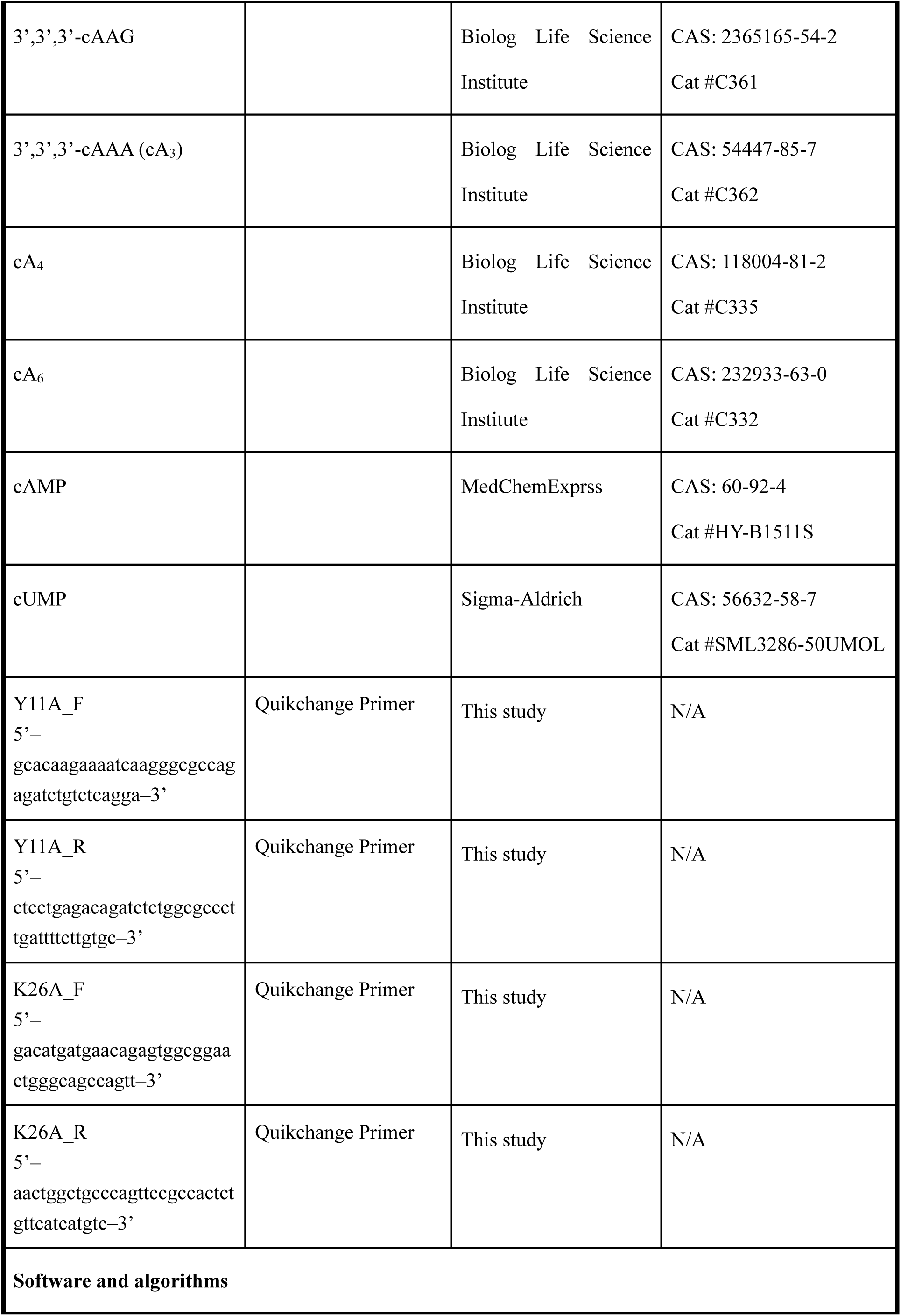
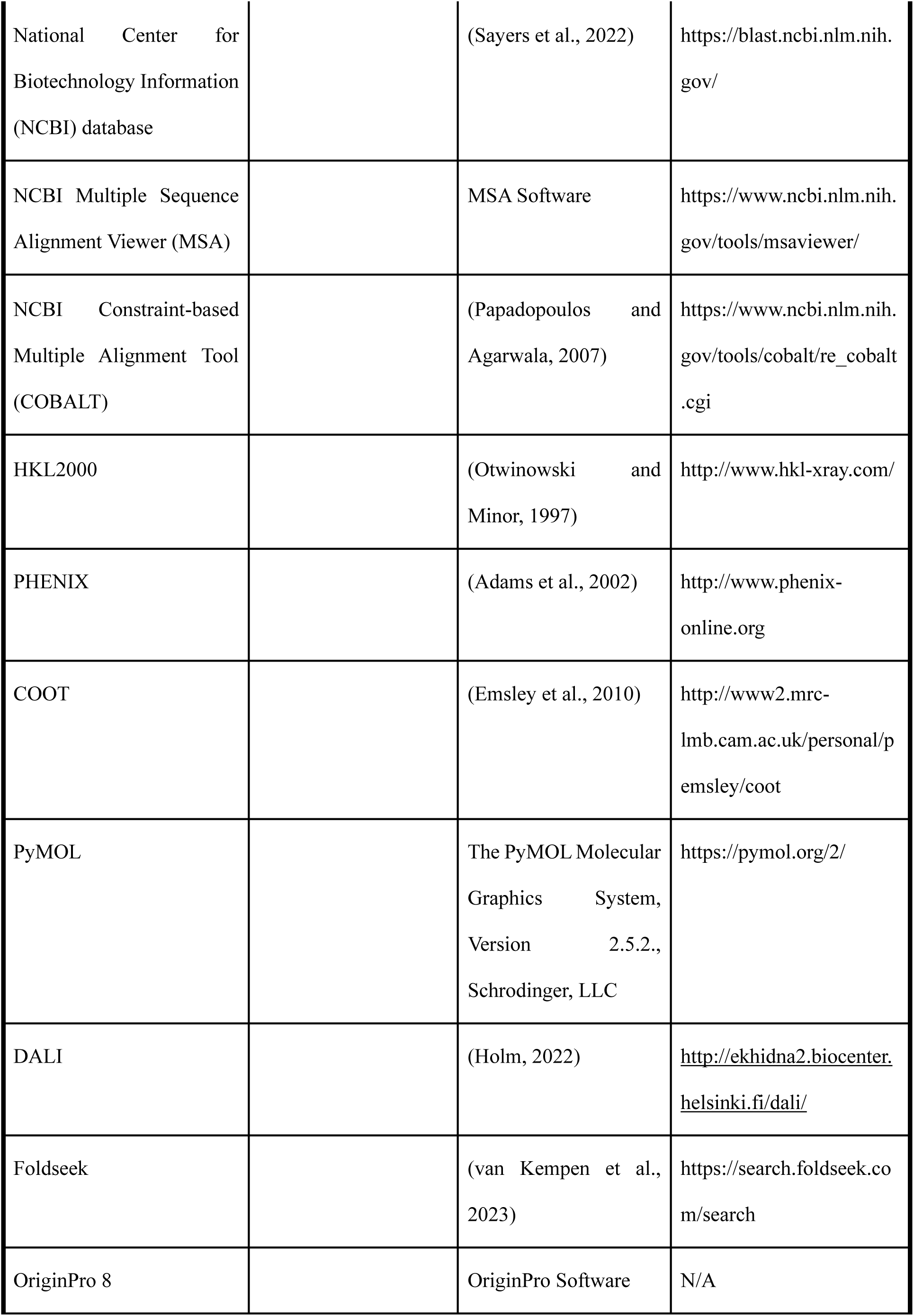
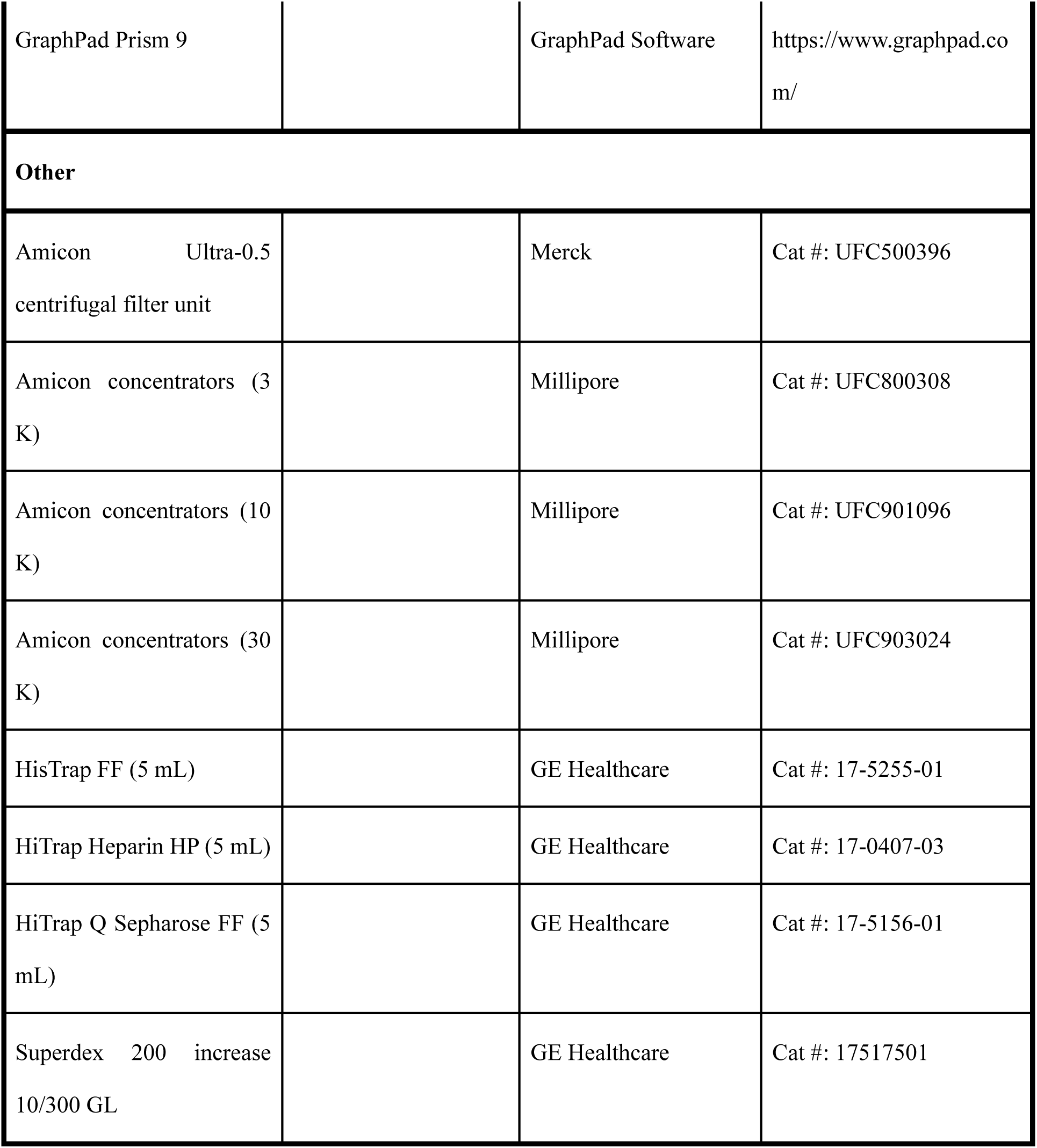

### RESOURCE AVAILABILITY

#### Materials Availability

All unique/stable reagents generated in this study are available from the Lead Contact with a completed Materials Transfer Agreement.

#### Data and Code Availability

The accession numbers for the coordinate and structure factors reported in this paper are PDB: 8IXZ (Acb2-3’,2’-cGAMP), 8J8O (Acb2-2’,3’-cGAMP), 8IY0 (Acb2-cA3), 8IY1 (Acb2-cAAG) and 8IY2 (Acb2-3’,3’-cGAMP-cA3). This paper does not report original code. Any additional information required to reanalyze the data reported in this paper is available from the lead contact upon request.

### EXPERIMENTAL MODEL AND SUBJECT DETAILS

#### Bacterial strains and phages

The bacterial strains and phages used in this study are listed in the Key Resources Table. The *P. aeruginosa* strains (BWHPSA011, ATCC 27853, PAO1) and *E. coli* strains (DH5ɑ) were grown in Lysogeny broth (LB) medium at 37°C both with aeration at 225 r.p.m. Plating was performed on LB solid agar with 10 mM MgSO4 when performing phage infections, and when indicated, gentamicin (50 µg ml^−1^ for *P. aeruginosa* and 15 µg ml^−1^ for *E. coli*) was used to maintain the pHERD30T plasmid. Gene expression was induced by the addition of L-arabinose (0.1%) unless stated otherwise. The *E. coli* BL21 (DE3) strain was used for recombinant protein overexpression and grown in Lysogeny broth (LB) medium. The cells were grown at 37°C until OD600nm reached 0.8 and then induced at 18°C for 12 h.

### METHOD DETAILS

#### Protein expression and purification

The PaMx33-Acb2, JBD67-Acb2, T4-Acb2, and CHI14-Acb2 genes were synthesized by GenScript. The full-length Acb2 gene was amplified by PCR and cloned into a modified pET28a vector in which the expressed Acb2 protein contains a His6-SUMO tag or His6 tag. The Acb2 mutants were generated by two-step PCR and were subcloned, overexpressed and purified in the same way as wild-type protein. The proteins were expressed in *E. coli* strain BL21 (DE3) and induced by 0.2 mM isopropyl-β-D-thiogalactopyranoside (IPTG) when the cell density reached an OD600nm of 0.8. After growth at 18°C for 12 h, the cells were harvested, re-suspended in lysis buffer (50 mM Tris–HCl pH 8.0, 300 mM NaCl, 10 mM imidazole and 1 mM PMSF) and lysed by sonication. The cell lysate was centrifuged at 20,000 g for 50 min at 4°C to remove cell debris. The supernatant was applied onto a self-packaged Ni-affinity column (2 mL Ni-NTA, Genscript) and contaminant proteins were removed with wash buffer (50 mM Tris pH 8.0, 300 mM NaCl, 30 mM imidazole). The fusion protein of Acb2 with His6-SUMO tag was then digested with Ulp1 at 18°C for 2 h, and then the Acb2 protein was eluted with wash buffer. The eluant of Acb2 was concentrated and further purified using a Superdex-200 increase 10/300 GL (GE Healthcare) column equilibrated with a buffer containing 10 mM Tris-HCl pH 8.0, 500 mM NaCl and 5 mM DTT. The purified protein was analyzed by SDS-PAGE. The fractions containing the target protein were pooled and concentrated.

The His6-Acb2 proteins bound to Ni-NTA beads were washed with wash buffer (50 mM Tris pH 8.0, 300 mM NaCl, 30 mM imidazole) and then eluted with the 50 mM Tris pH 8.0, 300 mM NaCl, 300 mM imidazole. The eluant of His6-Acb2 was concentrated, then further purified and analyzed as described above.

#### Crystallization, data collection and structural determination

The crystals of Acb2 were grown with reservoir solution containing 0.2 M Sodium bromide, 0.1 M Bis-Tris propane pH 6.5, 10% Ethylene glycol and 20% v/v PEG 3350 at 18°C. Prior to crystallization, cA3, cAAG, cA3+3’,3’-cGAMP or 3’,2’-cGAMP were mixed with the protein at a molar ratio of 2:1, respectively, and the concentration of Acb2 was 24 mg/mL. The crystals appeared overnight and grew to full size in about two to three days. The crystals were cryoprotected in the reservoir solution containing 20% glycerol before its transferring to liquid nitrogen.

All the data were collected at SSRF beamlines BL02U1 and BL19U1, integrated and scaled using the HKL2000 package (Otwinowski and Minor, 1997). The initial model of Acb2 was used from PDB: 8H2X. The structures of Acb2 and its complex with cyclic oligonucleotides were solved through molecular replacement and refined manually using COOT (Emsley et al., 2010). The structure was further refined with PHENIX (Adams et al., 2002) using non-crystallographic symmetry and stereochemistry information as restraints. The final structure was obtained through several rounds of refinement. Data collection and structure refinement statistics are summarized in Table 1.

#### Isothermal titration calorimetry binding assay

The dissociation constants of binding reactions of Acb2 or Acb2 mutants with the cA3/ cAAG/ 3’,2’-cGAMP/ 3’,3’-cGAMP/ 2’,3’-cGAMP/ c-di-AMP/ c-di-GMP/ 3’,3’-cUU/ 3’,3’-cUA/ 3’,3’-cUG/ cUMP/ cCMP/ cAMP/ cA6/ cA4 were determined by isothermal titration calorimetry (ITC) using a MicroCal ITC200 calorimeter. Both proteins and cyclic-oligonucleotides were desalted into the working buffer containing 20 mM HEPES pH 7.5 and 200 mM NaCl. The titration was carried out with 19 successive injections of 2 µL cA3/cAAG/cA6/cA4 at the 0.04 mM concentration, spaced 120 s apart, into the sample cell containing the Acb2 or Acb2 mutants with a concentration of 0.01 mM by 700 rpm at 25°C. Correspondingly, the 3’,2’-cGAMP/3’,3’-cGAMP/2’,3’-cGAMP/c-di-AMP/c-di-GMP/3’,3’-cUU/ 3’,3’-cUA/3’,3’-cUG/cUMP/cAMP at the 0.4 mM concentration was titrated into 0.1 mM Acb2 or Acb2 mutants at the same experimental conditions. The Origin software was used for baseline correction, integration, and curve fitting to a single site binding model.

#### Native-PAGE assay

Acb2 or Acb2 mutants was pre-incubated with cyclic-dinucleotides for 10 min at 18°C, where Acb2 or Acb2 mutants was 14.3 μM and the concentrations of cyclic-dinucleotides ranged from 1.8 to 7.2 μM (1.8, 3.6, 7.2 μM). Products of the reaction were analyzed using 5% native polyacrylamide gels and visualized by Coomassie blue staining.

#### High-performance liquid chromatography (HPLC)

40 μM Acb2 or Acb2 mutants was pre-incubated with 4 μM cA3 for 10 min at 18°C. Proteinase K was subsequently added to the reaction system at a final concentration of 0.5 μM and the reaction was performed at 58°C for 1 h. Reaction products were transferred to Amicon Ultra-15 Centrifugal Filter Unit 3 kDa and centrifuged at 4°C, 4,000 g. The products obtained by filtration were further filtered with a 0.22 μm filter and subsequently used for HPLC experiments. The HPLC analysis was performed on an Agilent 1200 system with a ZORBAX Bonus-RP column (4.6 × 150 mm). A mixture of acetonitrile (2%) and 0.1% trifluoroacetic acid solution in water (98%) were used as mobile phase with 0.8 mL/min. The compounds were detected at 254 nm.

#### *In vitro* NucC activity assay

For nuclease activity assay, a pUC19 plasmid was used as substrate. Pa-NucC (10 nM) and cA3 molecules (5 nM) were mixed with 0.5 μg DNA in a buffer containing 25 mM Tris-HCl pH 8.0, 10 mM NaCl, 10 mM MgCl2, and 2 mM DTT (20 μL reaction volume), incubated 10 min at 37°C, then separated on a 1% agarose gel. Gels were stained with Goldview and imaged by UV illumination.

To determine the function of Acb2, 50 nM Acb2 or its mutants were pre-incubated with the system at 18℃ for 15 min, and the subsequent reaction and detection method was as described above. To examine whether the released molecule from Acb2 is able to activate NucC, 5 nM cA3 was incubated with 50 nM Acb2 for 15 min at 18℃. Proteinase K was subsequently added to the reaction system at a final concentration of 1 μM and the reaction was performed at 58℃ for 1 h, then the proteinase K-treated samples were heated with 100℃ for 10 min to extinguish proteinase K and the subsequent detection method was as described above.

#### Episomal gene expression

The shuttle vector that replicates in *P. aeruginosa* and *E. coli*, pHERD30T (Qiu et al., 2008) was used for cloning and episomal expression of *P. aeruginosa* BWHPSA011 Type II-A or ATCC 27853 Type III-C CBASS operons into the PAO1 WT strain. This vector has an arabinose-inducible promoter and a selectable gentamicin marker. Vector was digested with SacI and PstI restriction enzymes and then purified. Inserts were amplified by PCR using bacterial overnight culture or phage lysate as the DNA template, and joined into the pHERD30T vector at the SacI-PstI restriction enzyme cut sites by Hi-Fi DNA Gibson Assembly (NEB) following the manufacturer’s protocol. The resulting plasmids were transformed into *E. coli* DH5ɑ. All plasmid constructs were verified by sequencing using primers that annealed to sites outside the multiple cloning site. *P. aeruginosa* cells were electroporated with the pHERD30T constructs and selected on gentamicin.

#### Plaque assays

Plaque assays were conducted at 37°C with solid LB agar plates. 150 μL of overnight bacterial culture was mixed with top agar and plated. Phage lysates were diluted 10-fold then 2 μL spots were applied to the top agar after it had been poured and solidified.

#### Interferon reporter assay in human cell line

The PaMx33-Acb2 gene was codon-optimized for human expression and synthesized by GenScript with overhangs that enabled insertion into the XhoI–BamHI sites of pcDNA3 via Gibson assembly. The wild-type plasmid was modified by site-directed mutagenesis using the QuikChange protocol with the indicated primers, and Dpn1-digested to obtain all point mutants. All recombinant plasmids were transformed into XL1-Blue competent cells (Agilent) and sequenced for verification.

293T-Dual Null cells were cultured in DMEM (Gibco) supplemented with 10% FBS (Atlanta Biologics) (v/v) and 100 U/mL penicillin-streptomycin (Gibco) and maintained in 37°C incubators with 5% CO2. Four days prior to measurement, 293T-Dual cells were passaged and plated in 12-well tissue cultured-treated plates at 100000 cells/well. After 20 hours they were transfected with 100 ng of pcDNA3-hSTING and 100 ng of pcDNA3 empty vector or containing Acb2 using Fugene6 transfection reagent (Promega) according to its associated protocol. After 20 hours, the growth media was replaced with fresh growth media containing 50 µM 2’,3’-cGAMP or regular growth media as negative controls. The cells were further incubated for 18 h, and the media was harvested to measure luciferase activity using the QuantiLuc system (Invivogen). The cells were directly lysed in 1 × LSB and western blot was performed on cell lysates to verify expression of STING and Acb2.

#### Western blot

A rabbit Acb2 polyclonal antibody was generated by a commercial vender (GenScript) using a synthetic peptide from Acb2 (CHNRDEITRIANAEP). The polyclonal Acb2 antibody was further purified by antigen affinity (GenScript).

After harvesting conditioned media, cells were directly lysed on the plate using 1× LSB. Lysates were separated on a SurePage Bis-Tris polyacrylamide gel (GenScript) and transferred to a nitrocellulose membrane using the semi-dry iBlot2 system (Invitrogen). The membrane was blocked for 1 h at room temperature (Intercept blocking buffer, Li-COR Biosciences), and incubated with primary antibodies (rabbit anti-STING (Cell Signaling Technologies), mouse anti-alpha-tubulin (Sigma-Aldrich), rabbit anti-Acb2 (GenScript)) overnight at 4°C. Following three washes in 1×TBS-0.1% tween, secondary antibody (Anti-rabbit or anti-mouse (Li-Cor Biosciences)) was added for 1 hour at room temperature, followed by three additional washes in TBS-T. Blots were imaged in IR using a Li-Cor Odyssey Blot Imager.

