## Supplementary figures for "Phage anti-CBASS protein simultaneously sequesters cyclic trinucleotides and dinucleotides"

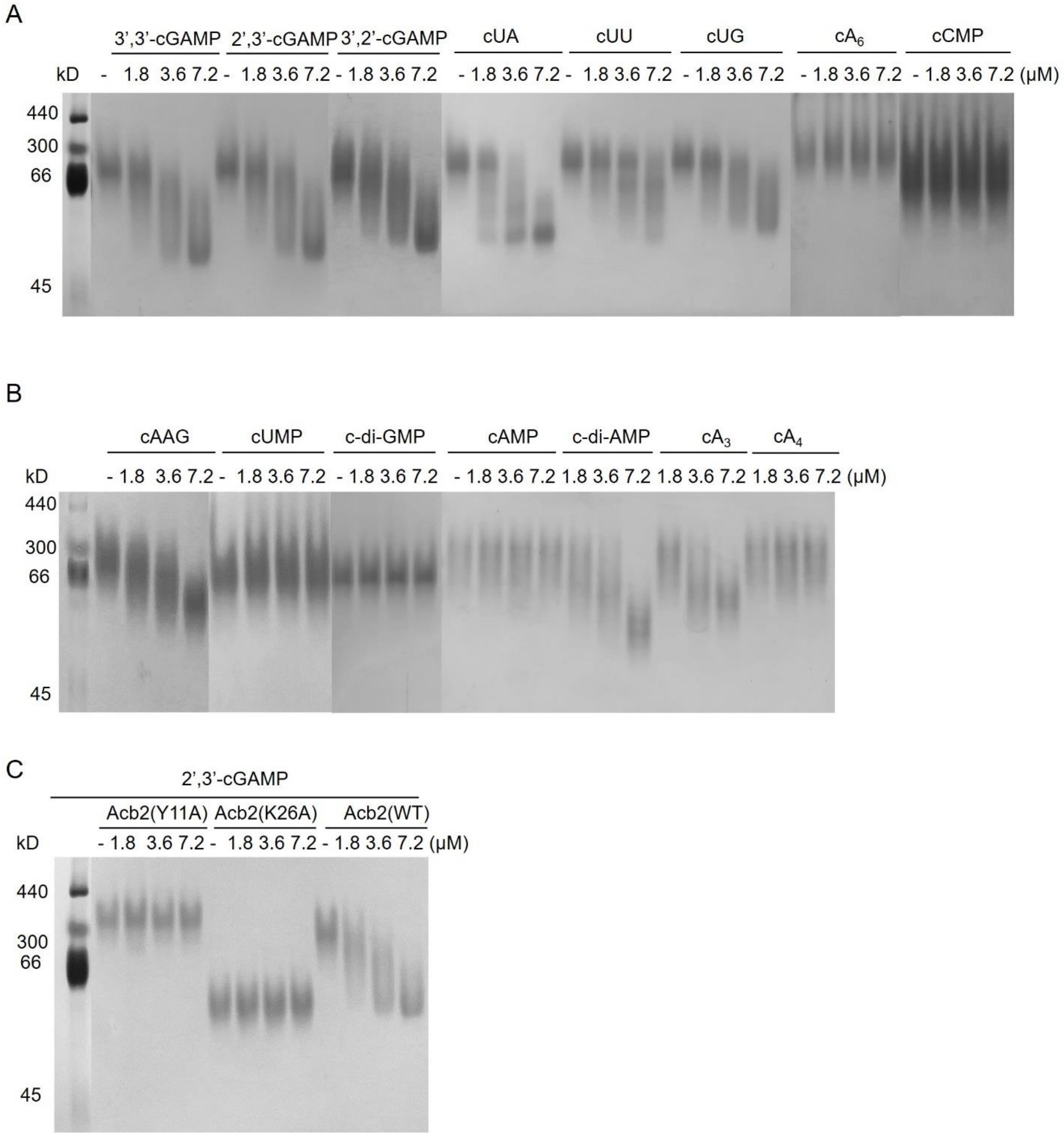


**Figure S1 The cyclic nucleotide binding spectrum of PaMx33-Acb2, related to Figure 1**

(A-C) Native PAGE assay showed the binding of PaMx33-Acb2 and its mutants to cyclic nucleotides.

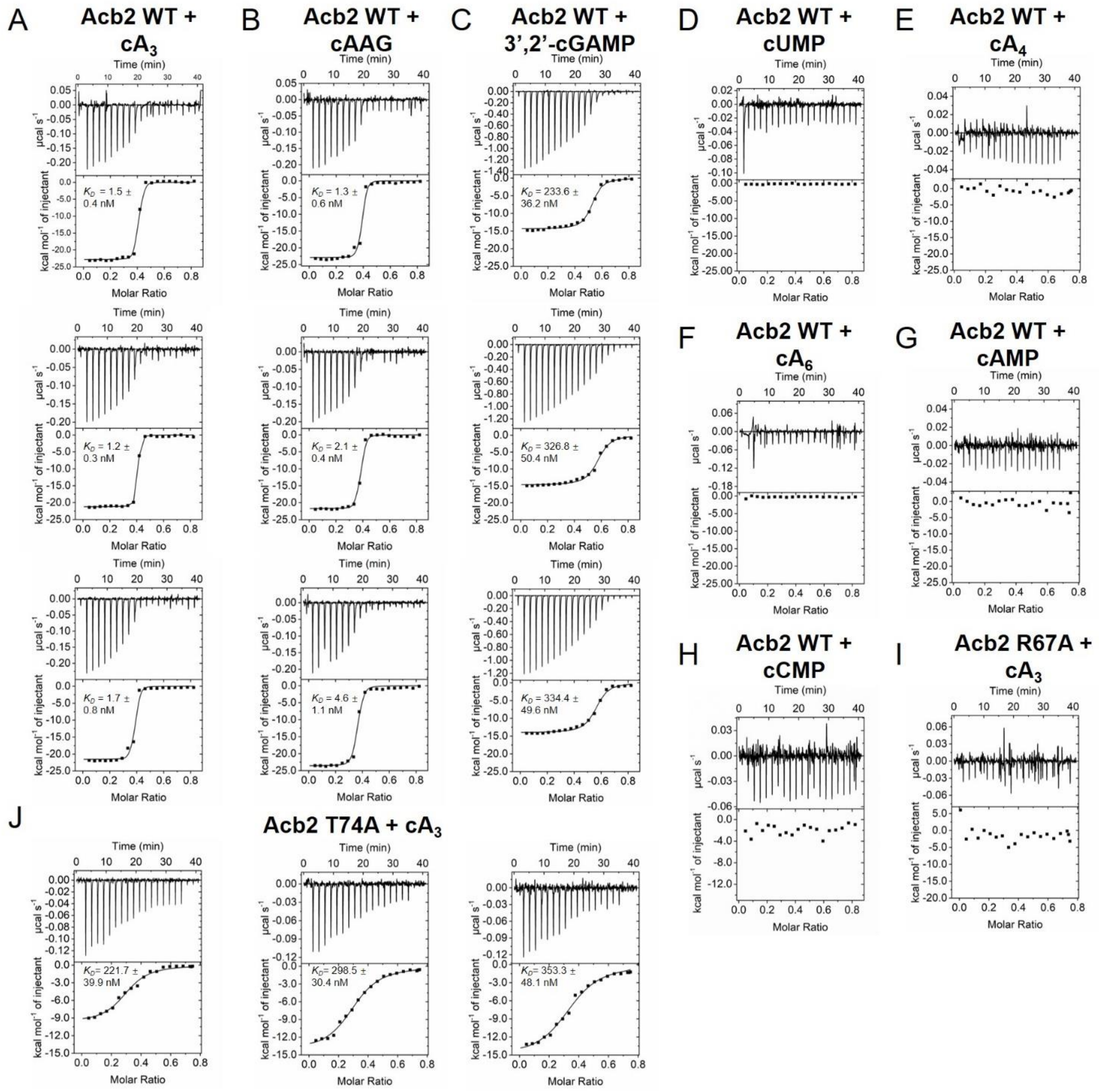


**Figure S2 The binding spectrum of PaMx33-Acb2 studied by ITC assays, related to Figure 1**

(A-H) ITC assays to test binding of PaMx33-Acb2 to cyclic nucleotides.

(I-J) ITC assays to test binding of $cA_3$ to PaMx33-Acb2 T74A and PaMx33-Acb2 R67A.

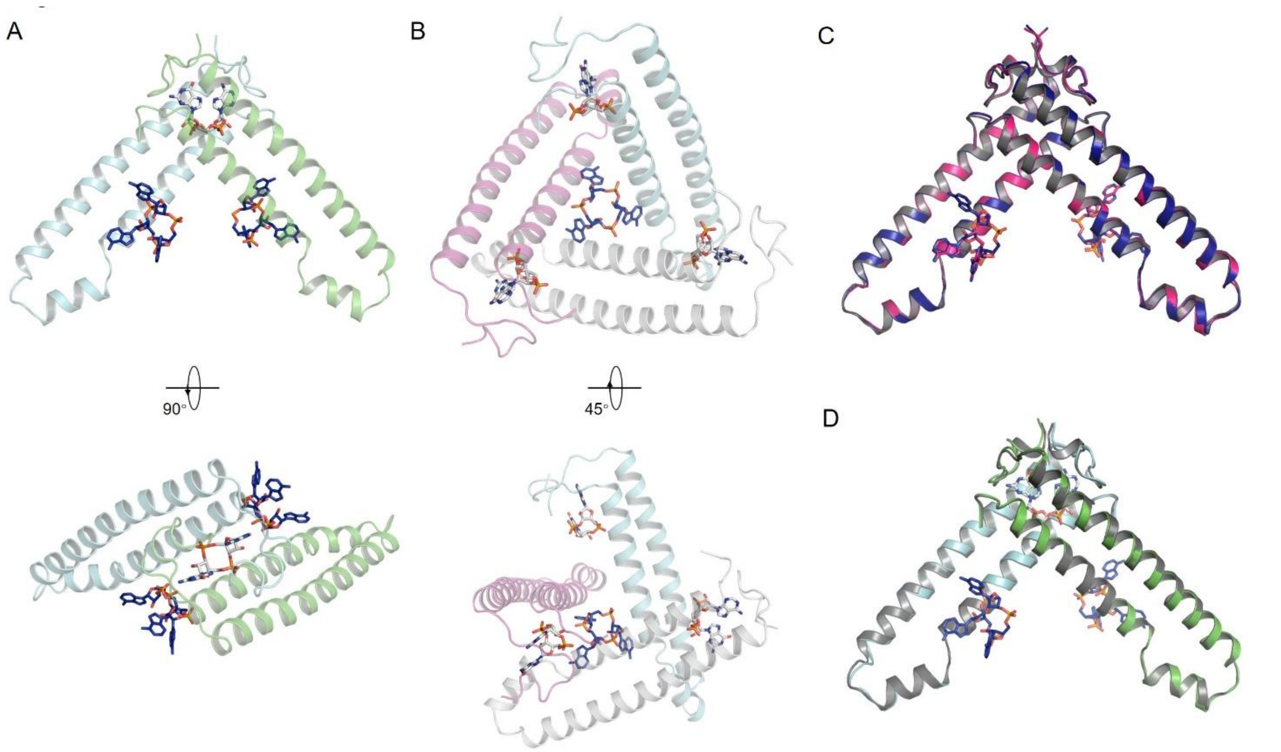


**Figure S3 The binding sites of cyclic trinucleotides and dinucleotides are different, related to Figure 2**

(A) An Acb2 dimer and the bound 3’,3’-cGAMP within the Acb2-3’,3’-cGAMP structure is shown. The Acb2-$cA_3$ structure is aligned to the structure and only the two $cA_3$ molecules are shown.

(B) An Acb2 trimer and the bound $cA_3$ within the Acb2-$cA_3$ structure is shown. The Acb2-3’,3’-cGAMP structure is aligned to the structure and only the three 3’,3’-cGAMP molecules are shown.

(C) Structural superimposition among Acb2-$cA_3$ (colored deep blue), Acb2-cAAG (colored hot pink) and apo Acb2 (colored grey) structures. Only an Acb2 dimer and the cyclic trinucleotides it is binding are shown.

(D) Structural superimposition between Acb2-$cA_3$-3’,3’-cGAMP and apo Acb2 structures. Only an Acb2 dimer and the cyclic nucleotides it is binding are shown.

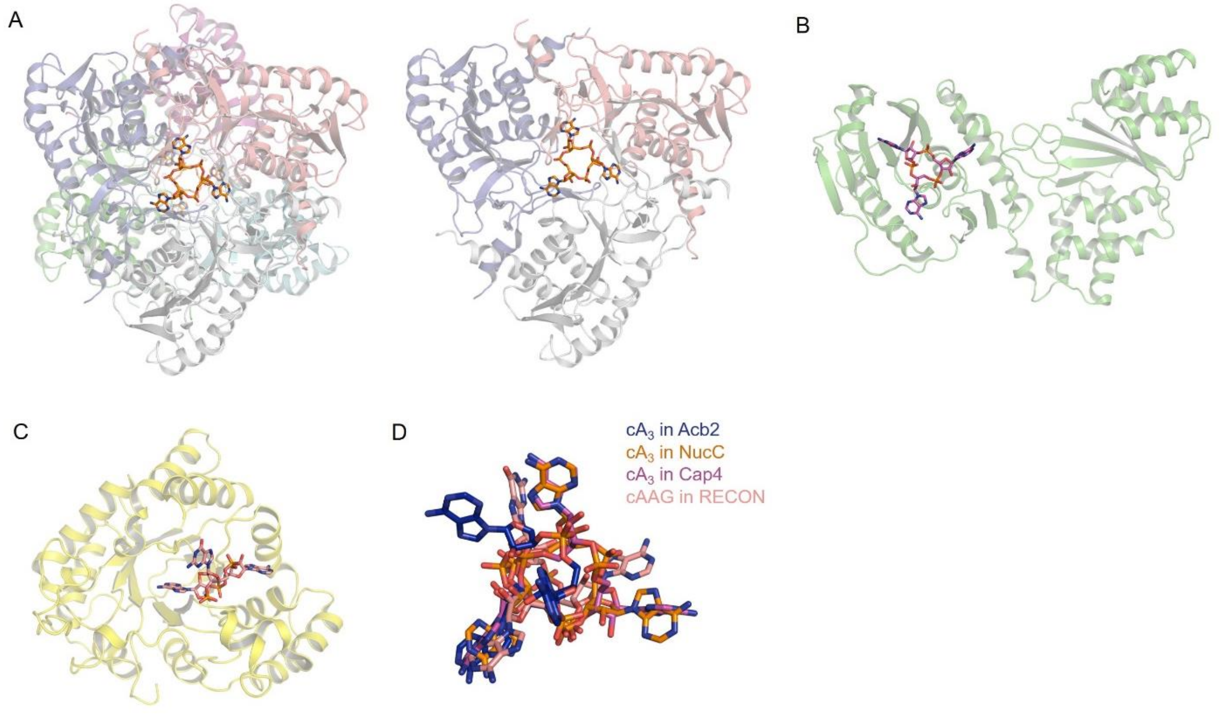


**Figure S4 Acb2 binds to $cA_3$ with a novel fold, related to Figure 3**

(A) The binding of $cA_3$ in NucC from *Escherichia coli* (PDB code: 6P1H). A NucC hexamer bound with two $cA_3$ molecules and a NucC trimer bound with one $cA_3$ molecule are shown in the left and right, respectively.

(B) The binding of $cA_3$ in Cap4 from *Acinetobacter baumannii* (PDB code: 6WAN).

(C) The binding of cAAG in RECON from *Mus musculus* (PDB code: 6M7K).

(D) $cA_3$ and cAAG molecules in the structures of Acb2-$cA_3$, NucC-$cA_3$, Cap-$cA_3$ and RECON-cAAG are aligned together and highlighted.

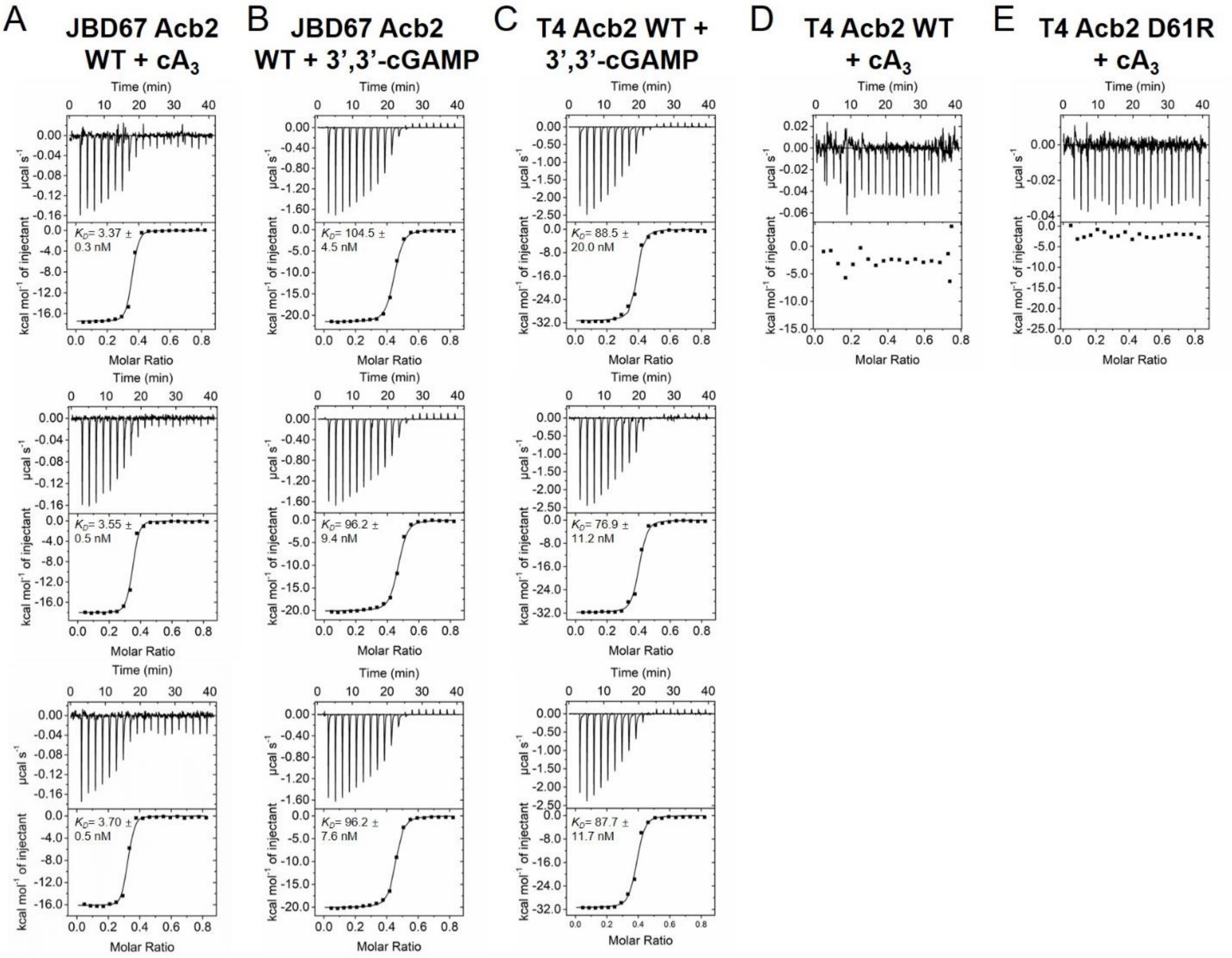


**Figure S5 The binding spectrums of JBD67-Acb2 and T4-Acb2 studied by ITC assays, related to Figure 4**

(A-B) ITC assays to test binding of cyclic oligonucleotides to JBD67-Acb2.

(C-D) ITC assays to test binding of 3',3'-cGAMP to T4-Acb2.

(E) ITC assays to test binding of $cA_3$ to T4-Acb2 D61R.

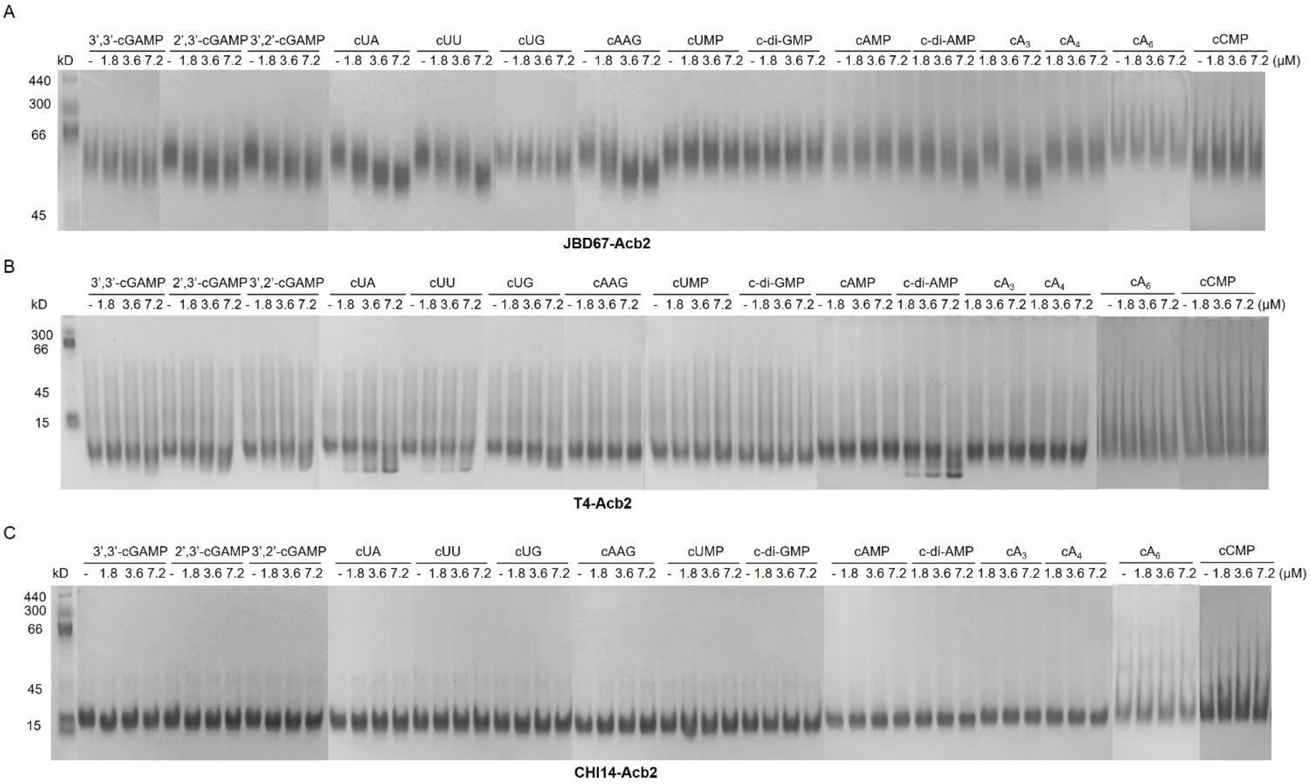


**Figure S6 The binding spectrums of Acb2 homologs studied by native PAGE, related to Figure 4**
(A-C) Native PAGE assay showed the binding of cyclic nucleotides to JBD67-Acb2, T4-Acb2 and CHI14-Acb2. The proteins were incubated with small molecules at indicated concentrations. Then the samples were subjected to native PAGE.

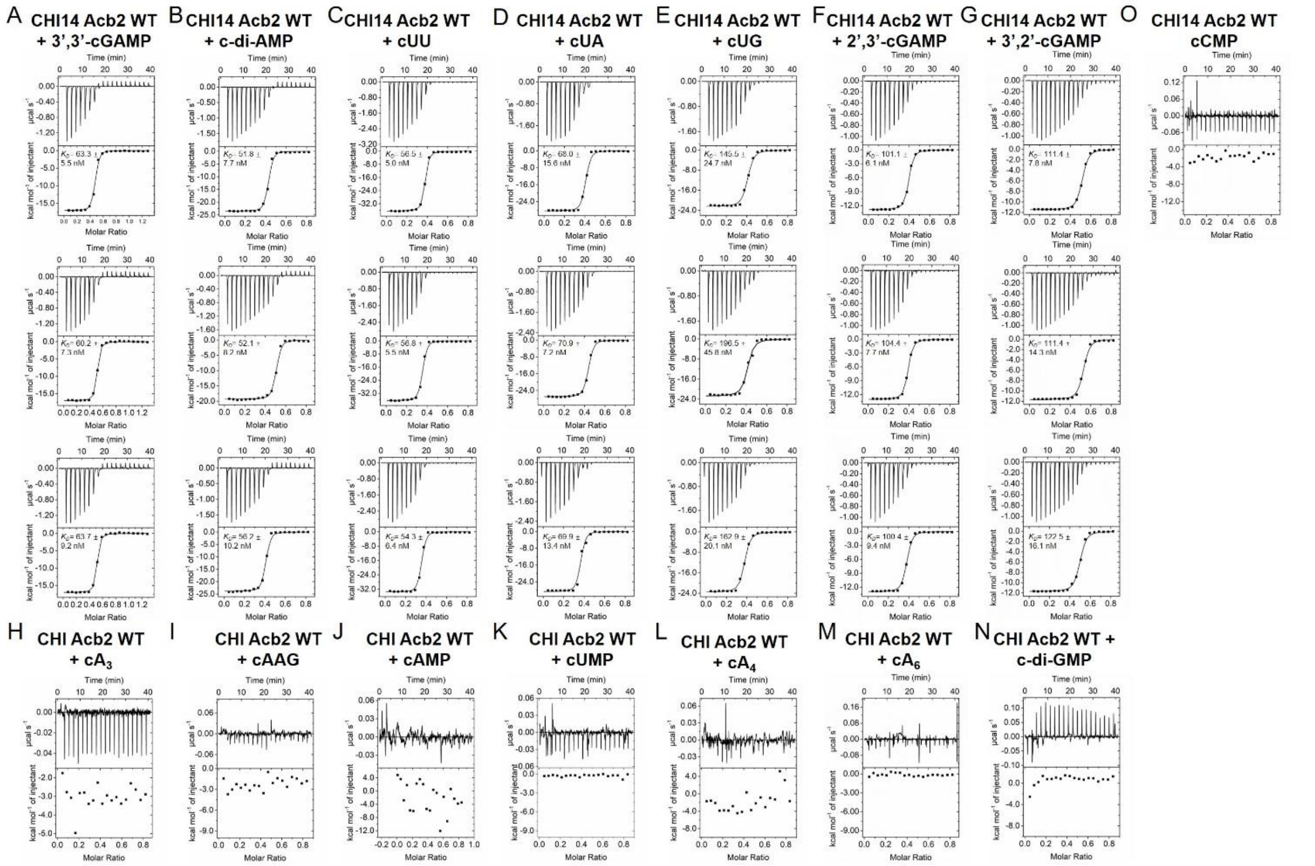


**Figure S7 The binding spectrum of CHI14-Acb2 studied by ITC assays, related to Figure 4**

(A-O) ITC assays to test binding of cyclic nucleotides to CHI14-Acb2.
